# *Salmonella* succinate utilisation is inhibited by multiple regulatory systems

**DOI:** 10.1101/2022.12.21.521472

**Authors:** Nicolas Wenner, Xiaojun Zhu, Will P. M. Rowe, Kristian Händler, Jay C. D. Hinton

## Abstract

Succinate is a potent immune signalling molecule that is present in the mammalian gut and within macrophages. Both of these niches are colonised by the pathogenic bacterium *Salmonella enterica* serovar Typhimurium during infection. Succinate is a C_4_-dicarboyxlate that can serve as a source of carbon for bacteria. When succinate is provided as the sole carbon source for *in vitro* cultivation, *Salmonella* and other enteric bacteria exhibit a slow growth rate and a long lag phase. This growth inhibition phenomenon was known to involve the sigma factor RpoS, but the genetic basis of the repression of bacterial succinate utilisation was poorly understood. Here, we used an experimental evolution approach to isolate fast-growing mutants during growth of *S*. Typhimurium on succinate containing minimal medium.

Our approach reveals novel RpoS-independent systems that inhibit succinate utilisation. The CspC RNA binding protein restricts succinate utilisation, an inhibition that is antagonised by high levels of the small regulatory RNA (sRNA) OxyS. We discovered that the Fe-S cluster regulatory protein IscR inhibits succinate utilisation by repressing the C_4_-dicarboyxlate transporter DctA.

The RNA chaperone Hfq, the exoribonuclease PNPase and their cognate sRNAs function together to repress succinate utilisation *via* RpoS induction. Furthermore, the ribose operon repressor RbsR is required for the complete RpoS-driven repression of succinate utilisation, suggesting a novel mechanism of RpoS regulation.

Our discoveries shed light on redundant regulatory systems that tightly regulate the utilisation of succinate. We propose that the control of central carbon metabolism by multiple regulatory systems in *Salmonella* governs the infection niche-specific utilisation of succinate.

## Introduction

Metabolic versatility is a key property that allows pathogenic enteric bacteria to thrive both during infection of mammals and in the wider environment [1]. C_4_-dicarboyxlates are an important part of the bacterial catabolic repertoire, which can be utilised as a sole carbon and energy source (C-source) [2]. In the mammalian gut, the C_4_-dicarboyxlate succinate is an abundant C-source that is provided by the microbiota in response to the presence of dietary fibre [3]. *Salmonella enterica* serovar Typhimurium (*S.* Typhimurium) is one of the best understood enteropathogenic bacterium [4] which efficiently catabolises succinate during intestinal colonisation to enhance the growth [5].

As well as colonising the mammalian gut, *Salmonella* can also cross the intestinal epithelial barrier and invade several types of tissues [6]. An important element of the pathogenic lifestyle of *S.* Typhimurium involves the hijacking of macrophages, and the intracellular proliferation of the bacteria within *Salmonella*-containing vacuoles (SCVs) [7]. Recently, it has been discovered that macrophages undergo metabolic reprogramming during bacterial infection, leading to the build-up of tricarboxylic acid (TCA) cycle intermediates, including succinate [8]. This C_4_-dicarboxylate acts as an important proinflammatory molecule that is also involved in hypoxic and metabolic signalling [9, 10]. Following infection by *S.* Typhimurium, high levels of succinate accumulate within macrophages [11]. However, *Salmonella* does not use this succinate to fuel growth as glucose and the glycolytic intermediate 3-phosphoglycerate are the key intra-macrophage C-sources [11–13]

The inactivation of key succinate catabolic genes does not reduce the ability of *S.* Typhimurium to replicate in murine macrophages, but stimulates intracellular proliferation [14]. Although succinate is not utilised as a C-source by *Salmonella* in the SCV, the metabolite does act as a crucial signal molecule for the induction of the *Salmonella* Pathogenicity Island 2 (SPI2) system [13], which is required for macrophage infection [15].

During *in vitro* cultivation, *S.* Typhimurium exhibits a particularly extended lag phase in minimal axenic media containing succinate as sole C-source; in contrast, succinate supports the rapid growth of other enteric bacteria such as *Escherichia coli* [16], *via* the succinate dehydrogenase (SDH) multi-enzyme complex that oxidises succinate into fumarate [17]. Subsequent, bacterial replication with succinate involves the generation of all cellular components *via* gluconeogenesis [2, 18].

The stress response sigma factor σ^38^ (RpoS) is a global transcriptional regulator that modulates diverse facets of *Salmonella* biology including stress-resistance, immobilised growth, virulence and nutrient assimilation [19–22]. RpoS inhibits *in vitro* growth upon succinate by repressing transcription of the *sdhCDAB* operon (*sdh*) and other TCA cycle genes [23–25].

Because *Salmonella* utilises succinate for colonisation of the inflamed gut [5] but not for intra-macrophage proliferation [13, 14], we hypothesised that *Salmonella* had evolved multiple genetic regulatory mechanisms to tightly control the niche-dependent utilisation of this infection-relevant molecule.

Here, we devised an *in vitro* experimental strategy to search for novel regulatory mechanisms involved in the modulation of succinate utilisation. Our genetic dissection identified two novel RpoS-independent regulatory mechanisms that repress succinate utilisation *via* the CspC and IscR regulatory proteins. In addition, the modulation of RpoS activity by Hfq, PNPase and RbsR also impacted upon succinate utilisation. We propose that this multi-factorial system ensures that succinate is only catabolised at the right place and at the right time during infection to permit effective niche adaptation.

## Results and Discussion

### Growth inhibition and evolution in succinate minimal medium

Enteric bacteria possess the catabolic enzymes and efficient uptake systems required to grow with C_4_-dicarboxylates as sole C-source [2, 26]. However, some environmental and clinical isolates of *Escherichia coli* and *Salmonella* have surprisingly slow growth rates in succinate-containing minimal media [27–29]. To investigate this phenomenon in pathogenic and non-pathogenic enteric bacteria, we assessed the growth of four bacterial species on agar plates containing succinate as a sole C-source (M9+Succ). We studied growth for up to 96 hours, and used a variety of laboratory strains with an emphasis on *Salmonella* (Fig 1).

**Figure 1:**
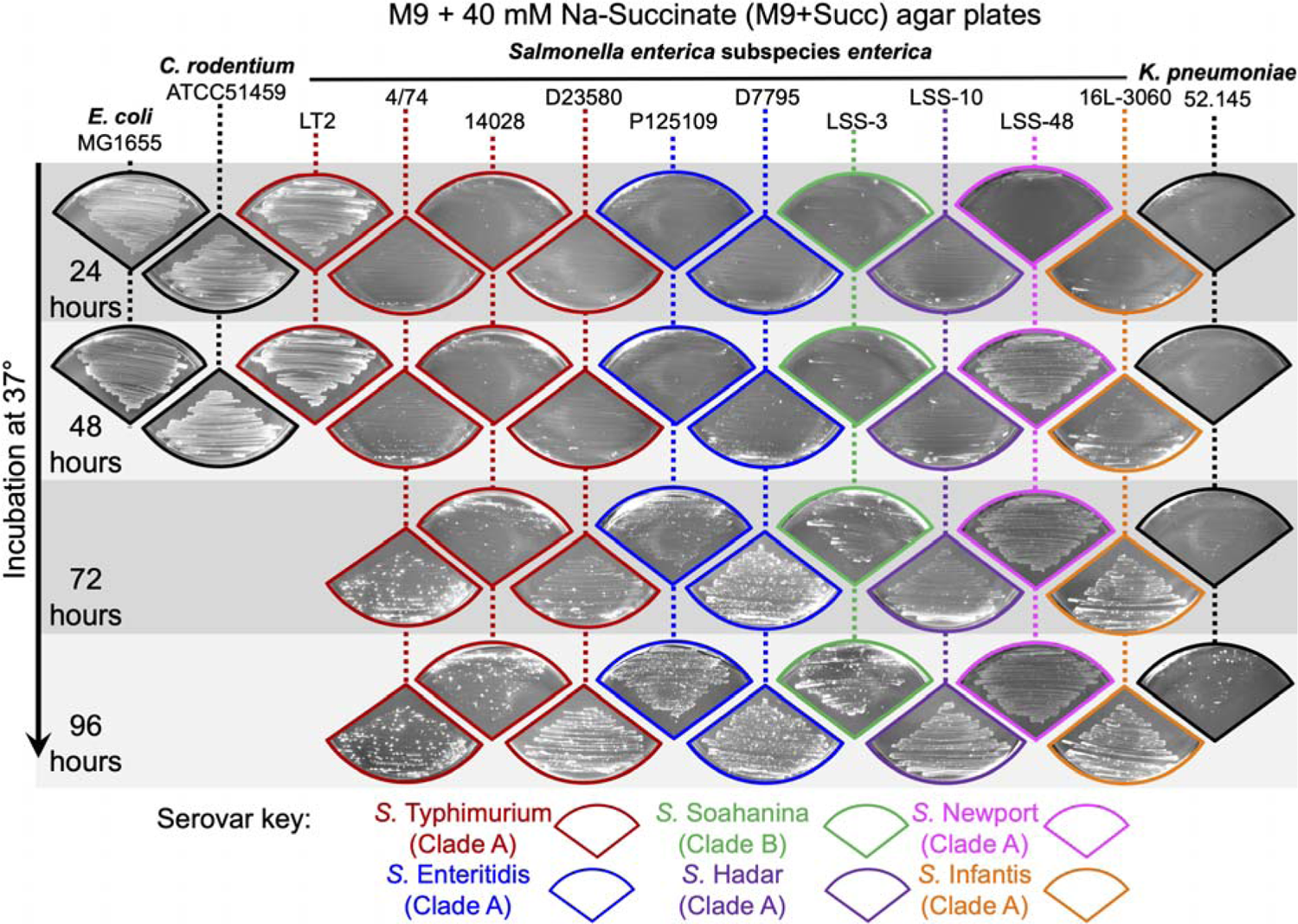
Inhibition of growth of most *Salmonella* serovars and *K. pneumoniae* on succinate minimal medium. The indicated strains of *E. coli*, *Salmonella*, *Citrobacter* and *Klebsiella* were spread on M9+Succ agar plates and incubated at 37°C. Photographs of bacterial growth were taken every 24 hours for 2 days or 4 days. For *Salmonella enterica* isolates, the serovar and the clade A/B status [33, 34] are indicated by the colour of the picture frame. Experiments were carried out as biological triplicates, and a representative picture is shown for each strain.

For *S. enterica* serovars Typhimurium and Enteritidis, we tested the growth of the well-characterised *S.* Typhimurium strains LT2, 4/74 and 14028 and of the *S.* Enteritidis strain P125109. In addition, we assessed the growth of multidrug resistant *S.* Typhimurium ST313 strain D23580 and *S.* Enteritidis strain D7795, two representative strains that cause invasive non-typhoidal *Salmonella* disease in Africa [30–32]. We included the reptile-associated *Salmonella* serovars Soahanina, Hadar, Newport and Infantis which belong to the two metabolically-distinct clades A and B of *S. enterica* [33, 34]. We and others have previously generated genome sequences of most of the *Salmonella* strains that were used [32,33,35]. We also tested a multidrug resistant strain of *Klebsiella pneumoniae* (strain KP52.145) [36], a *Citrobacter rodentium* strain (ATCC51459) [37] and the classic laboratory strain *E. coli* K-12 strain MG1655.

After 24 hours of incubation at 37°C, the only strains that displayed substantial growth with succinate as a C-source were *S.* Typhimurium LT2, *E. coli* MG1655 and *C. rodentium* ATCC51459. Following 2-3 days of incubation, large colonies were observed within the bacterial lawns of the *K. pneumoniae* strain and the other *Salmonella* isolates. The only *Salmonella* serovar that displayed substantial growth after 2 days was *S.* Newport, showing that growth on succinate is a serovar-dependent phenotype.

Previous experiments in liquid minimal medium containing succinate as sole C-source showed that *S.* Typhimurium exhibited a particularly long lag phase [16, 23]. This extended lag time could reflect a particularly slow metabolic remodelling, preparing *Salmonella* for the exponential phase in the presence of succinate. Alternatively, robust inhibition of succinate assimilation might be occurring under these conditions, preventing growth until spontaneous fast-growing mutants (hereafter referred as Succ^+^ mutants) have emerged.

To test these hypotheses, we assessed the growth of the well-characterised *S.* Typhimurium strain 4/74 (henceforth referred to as 4/74 or *Salmonella*) in liquid M9+Succ media inoculated with a stationary phase culture made in rich medium (LB). The four independent 4/74 cultures (cultures I-IV) exhibited the reported 30-35 hour lag time at 37°C [16, 23] (Fig 2A). We collected the *Salmonella* that eventually reached stationary phase and cultured the bacteria in LB for two passages before re-inoculation in M9+Succ. For all the succinate-evolved cultures, the lag time in M9+Succ was reduced to 4-5 hours, and stationary phase was reached after approximately 14 hours (Fig 2B).

**Figure 2.**
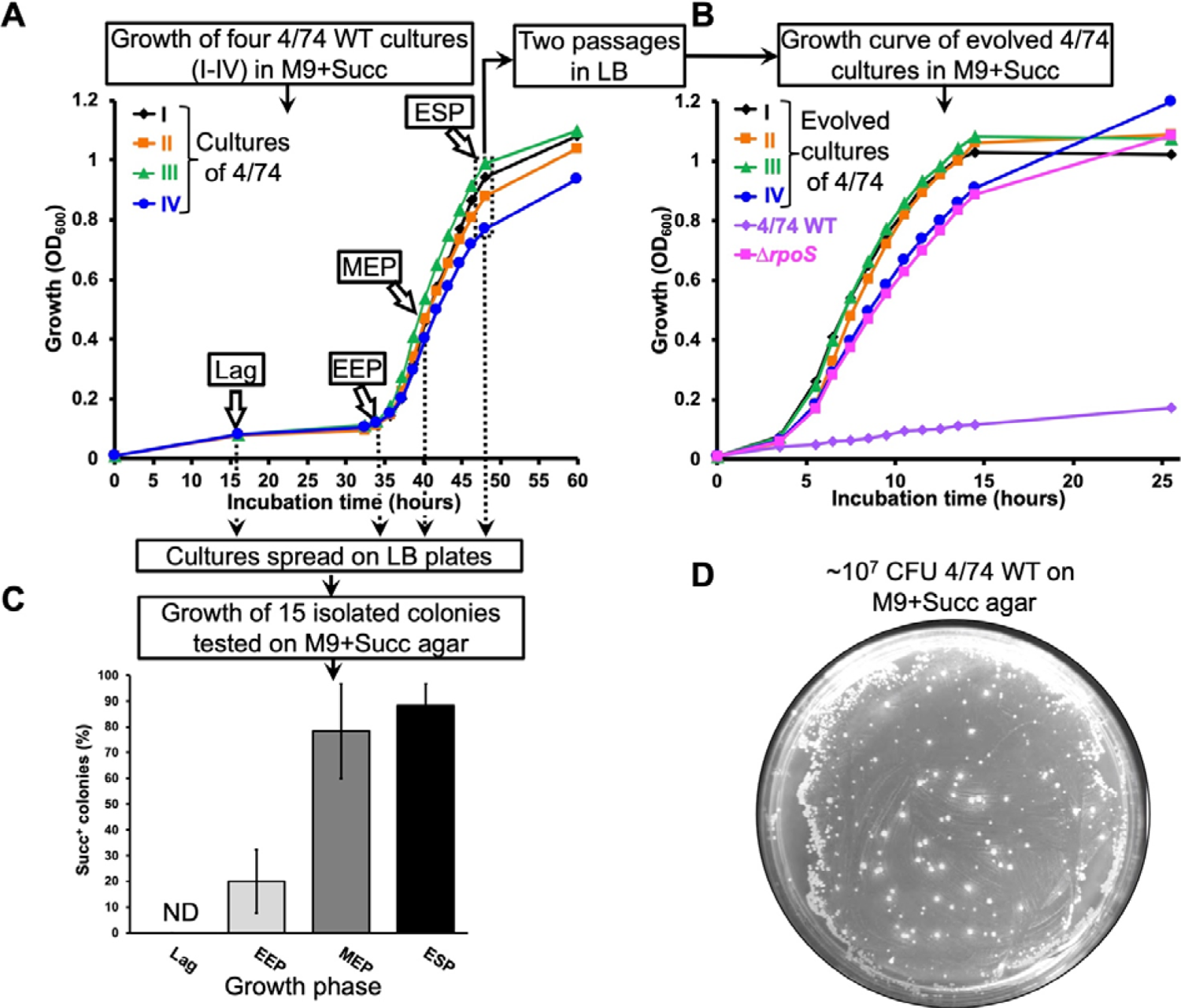
Experimental evolution of *S.* Typhimurium in succinate minimal medium. (**A**) Growth of *S.* Typhimurium 4/74 displays an extended lag time in M9+Succ medium. The growth curves of four independent cultures (I-IV) of 4/74 WT in M9+Succ are presented. Cultures were inoculated with bacteria grown beforehand to stationary phase in LB. the succinate evolved bacteria were harvested in ESP (50 µL of culture) and were grown twice in LB prior to re-inoculation in M9+Succ. (**B**) The succinate-evolved bacteria grown fast in M9+Succ in comparison with the WT strain. Growth curves of the succinate evolved cultures I-IV from (**A**) in M9+Succ are presented: the 4/74 WT and Δ*rpoS* (JH3674) strains were included as controls. (**C**) Succinate fast growing (Succ^+^) mutants were detected in liquid M9+Succ 4/74 cultures. Cultures from (**A**) were spread on LB plates at the indicated growth phase and growth with succinate was assessed for 15 isolated colonies *per* replicate on M9+Succ agar plates after 48 hours of incubation. The graph shows the proportion (%) of Succ^+^ clones. ND= not detected. (**D**) Succ^+^ spontaneous mutants emerge from 4/74 WT bacterial lawns on M9+Succ agar plates. 4/74 WT cultures (∼10^7^ CFU) were spread on a M9+Succ agar plates and the picture of a representative plate was taken after 3 days of incubation at 37°C. For the growth curves (**A**&**B**), bacteria were grown at 37°C with aeration in 25 ml of M9+Succ (in 250 ml conical flasks) with an initial inoculum of ∼10^7^ CFU/mL (OD_600_=0.01). Growth phases are indicated in (**A**&**C**): Lag phase (Lag); Early exponential phase (EEP); Mid-exponential phase (MEP); Early stationary phase (ESP).

To investigate heritability of the succinate growth phenotype, the initial M9+Succ cultures (Fig 2A) were spread on LB agar plates at different stages of growth, and isolated colonies were tested on M9+Succ plates. Of 60 colonies obtained from lag phase, none grew faster than the wild type (Fig 2C). However, fast growing Succ^+^ mutants harvested from early exponential, mid-exponential and early stationary phase culture were detected at a frequency of 20 %, 78 % and 90 %, respectively (Fig 2C). When ∼10^7^ Colony Forming Units (CFU) of 4/74 wild-type (WT) were spread on M9+Succ plates, between 100 and 1000 Succ^+^ colonies grew in the bacterial lawn after 3 days of incubation (Fig 2D).

Collectively, these results indicated that, in our experimental setup, *Salmonella* growth upon succinate was consistently inhibited. The eventual initiation of exponential phase did not result from an orchestrated metabolic switch, but reflected the emergence of spontaneous mutants that efficiently utilised succinate, and proliferated to outcompete the WT bacteria. This Succ^+^ phenotype remained stable after two passages on LB medium, indicating that the trait was not a phase-variable phenomenon caused by epigenetic mechanisms, as has been observed for other reversible phenotypes in *Salmonella* [38, 39]. Moreover, our data suggest that other pathogenic bacteria, such as *Klebsiella* also suppress succinate assimilation (Fig 1).

Here, we define “succinate utilisation” as the ability of *Salmonella* to grow with succinate as a sole carbon and energy source. We selected *S.* Typhimurium strain 4/74 for further study of the suppression of succinate utilisation because it is the parent of strain SL1344, which has been used for a plethora of regulatory and infection studies in the past [40]. We aimed to identify novel genetic determinants involved in the control of the uptake and catabolism of this infection-relevant C-source.

### Identification of novel mutations that abolish inhibition of succinate utilisation

To identify mutations that ablate the inhibition of succinate utilisation, we used three complementary unbiased approaches. We first screened a collection of published *S.* Typhimurium 4/74 mutants [41] that lacked key regulatory proteins (Fig S1A), and focused on mutations that both promoted growth on succinate agar plates and in liquid medium with aeration. This screen revealed that mutants lacking the RNA chaperone Hfq (Δ*hfq*) and the polynucleotide phosphorylase (PNPase, mutant Δ*pnp*) had a Succ^+^ phenotype. Complementation with low-copy plasmids carrying *hfq^+^* or *pnp^+^*, restored the Succ^-^ WT phenotype in the corresponding mutant (Fig S3C & D).

To explore metabolic suppression in more depth, we used global Tn*5* transposon mutagenesis to generate insertions that promoted growth on succinate (Methods). RpoS has a key role in the inhibition of succinate utilisation [27–29] and *rpoS* inactivation did cause the drastic shortening of the lag time of 4/74 in M9+Succ, similarly to the succinate evolved cultures (Fig 2B, Fig S1B, Fig S3B). Therefore, we developed a strategy to avoid the selection of Succ^+^ *rpoS* mutants by constructing a strain that carried two chromosomal copies of *rpoS* (4/74 *rpoS^2X^*; Fig S1B-C). Following Tn*5* mutagenesis of the *rpoS^2X^* strain, individual Succ^+^ Tn*5* mutants were isolated. The Tn*5* insertions were P22-transduced into 4/74 WT and the Succ^+^ phenotype of the transductants was confirmed (Table 1).

**Table 1:** Fifteen mutations that stimulated growth of *S.* Typhimurium upon succinate. The mutation coordinates and the annotations correspond to the *S.* Typhimurium 4/74 reference genome (GenBank: CP002487.1)[35]. For the Tn*5* insertions, the coordinates of the nucleotide after which the transposon was mapped is indicated. Transposon orientation is indicated as follows. >: the Tn*5 aph* gene (Km^R^) is encoded on the positive DNA strand of the 4/74 chromosome; <: *aph* is encoded on the negative strand. When insertions and mutations were mapped within the 5’ untranslated regions (5’-UTR) of a transcribed gene, the coordinates of upstream transcription start site (TSS) is indicated, according to the SalComMac transcriptomic database (http://bioinf.gen.tcd.ie/cgi-bin/salcom.pl?db=salcom_mac_HL [108, 133]. ^”^IGR” denotes intergenic regions, “SNP” single nucleotide polymorphisms, “Indel” insertions & deletion and “aSD” denotes the anti-Shine-Dalgarno (CCTCCTT) sequence of the 16S rRNAs.

| **Tn5 mutants** |  |  |
| --- | --- | --- |
| **coordinate** | **Locus affected and gene product** | **Tn5 orientation** |
| 436621 | Insertion in *iraP* (*yaiB*), anti-adapter protein IraP | < |
| 436616 | Insertion in *iraP* | < |
| 2007729 | Insertion in *fliD*, flagellar hook associated protein FliD | > |
| 2007415 | Insertion in *fliD* | > |
| 4118507 | Insertion in *rbsR*, ribose operon repressor RbsR | > |
| 1893137 | Insertion in *cspC*, cold shock-like protein CspC | > |
| 1893511 | Insertion in the *yobJ-cspC* operon 5'-UTR (*yobJ-cspC* TSS: coordinate 1893683) | > |
| 1893502 | Insertion in the *yobJ-cspC* operon 5'-UTR | < |
| **Spontaneous Succ<sup>+</sup> mutants** |  |  |
| **Mutation name** | **Locus affected and gene product** | **Mutation coordinates & characterisation** |
| *oxyR*<sup>mut</sup> | *oxyR*, encoding the regulatory protein sensor for oxidative stress OxyR | Indel, additional CTA (leucine) codon between coordinates 4365235-4365236 in *oxyR* |
| *iscR*<sup>STOP</sup> | *iscR* (*yfhP*), encoding the Fe-S cluster regulator protein IscR | SNP G→A coordinate 2680848, premature stop codon (Q<sup>99</sup>→STOP) in *IscR* |
| *dctA*<sup>mut1</sup> | 5'-UTR of *dctA*, encoding the aerobic C<sub>4</sub>-dicarboxylate transporter protein DctA | SNP T→C position 3812916 in *dctA* 5'-UTR (*dctA* TSS coordinate 3812935) |
| *dctA*<sup>mut2</sup> | 5'-UTR of *dctA*, encoding the aerobic C<sub>4</sub>-dicarboxylate transporter protein DctA | SNP G→A position 3812910 in *dctA* 5'-UTR |
| *rbsR*<sup>mut</sup> | 12 bp insertion in *rbsR*, ribose operon repressor RbsR | Indel: in frame insertion of 4 codons (TATCAGGCGCTA→YQAL) between codons 255 and 256 of *rbsR*, coordinates 4118620-4118621 |
| *rrsH*<sup>mut</sup> | *rrsH*, encoding the 16S ribosomal RNA RrsH | SNP T→A coordinate 290706 in the aSD<sup>d</sup> of *rrsH*: CCTCCTT→CCACCTT |
| *rrsA*<sup>mut</sup> | *rrsA*, encoding the 16S ribosomal RNA RrsA | SNP C→A coordinate 4219140 in the aSD<sup>d</sup> of *rrsA*: CCTCCTT→CATCCTT |

In parallel, we isolated spontaneous Succ^+^ mutants from either M9+Succ agar plates (Fig 2D) or from liquid cultures (Methods). We first verified the RpoS positive (*rpoS^+^*) status of each spontaneous Succ^+^ mutant (Methods), and then used whole genome-sequencing to identify relevant nucleotide changes, which were associated with seven genes (Table 1).

These complementary genetic screens identified Succ^+^ mutants that carried Tn*5* insertions in *iraP*, *cspC*, *rbsR* and *fliD* or in the 5’ untranslated region (5’-UTR) of the *yobJ-cspC* operon (Table 1). In addition, a nonsense spontaneous mutation in *iscR* (*yfhP*) and a spontaneous in-frame insertion of 4 codons in *rbsR* were identified in spontaneous Succ^+^ mutants (Table 1). To independently confirm the function of these genes, λ *red* recombination was used to generate Δ*iraP*, Δ *C*, Δ*rbsR* and Δ *scR* deletion mutants. Each of the four mutants had the Succ^+^ phenotype (Fig 3). We confirmed that the corresponding WT proteins could inhibit succinate utilisation by plasmid-borne complementation experiments (Fig S3 E-H).

**Figure 3.**
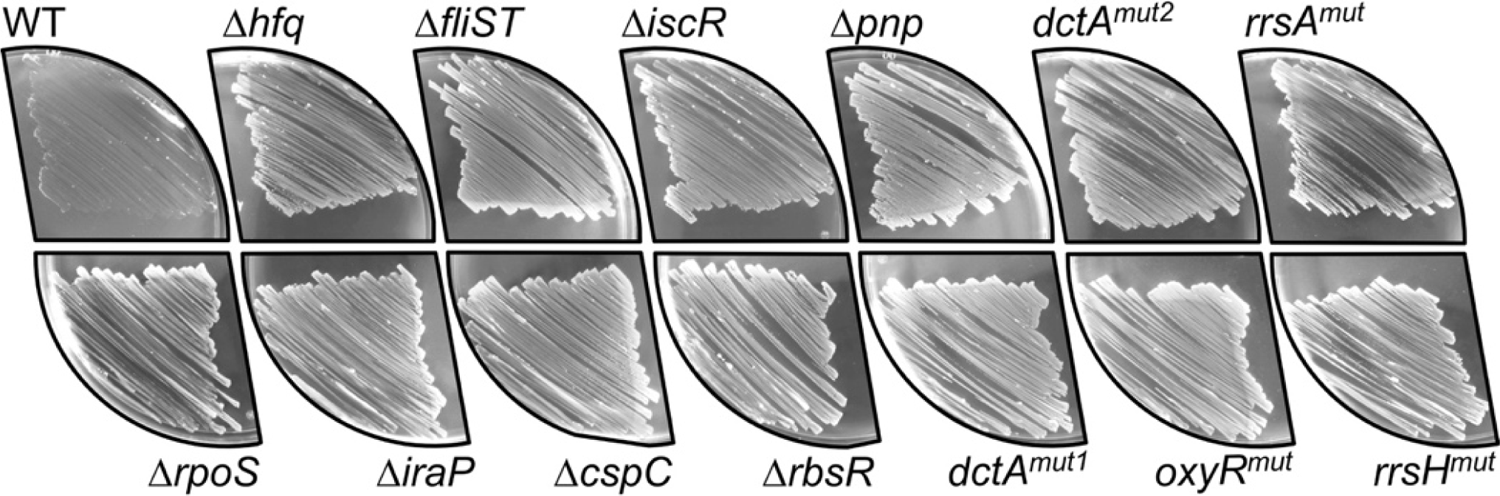
Growth phenotype of 5 genome-edited Succ^+^ mutants and 8 regulatory mutants on solid succinate minimal medium. *S*. Typhimurium strains 4/74 WT, Δ*rpoS* Δ*cspC* (SNW292), (SNW184), Δ*rbsR* (SNW294), Δ*pnp* (JH3649), *dctA^mut1^* (SNW160), *dctA^mut2^* (SNW315), *oxyR^mut^* (SNW318), *rrsA^mut^* (SNW336), *rrsH^mut1^* (SNW314), were spread on M9+Succ agar plates and incubated for 48 hours at 37°C. Experiments were carried out with biological triplicates and a representative picture is shown for each strain.

We found that inactivation of *fliD* alone did not cause a Succ^+^ phenotype (Fig S2A). The *fliD* gene is co-transcribed with the downstream *fliS* and *fliT* genes [42]. Tn*5* insertions may have polar effects on the expression of surrounding genes [43], raising the possibility that the *fliD*::Tn*5* insertion modulated expression of *fliS* or *fliT*. Our genetic dissection of the *fliDST* operon revealed that inactivation of either *fliS* or *fliT* caused the Succ^+^ phenotype, suggesting that the two genes contribute to the inhibition of succinate utilisation (Fig S2A). The Succ^+^ phenotype of the Δ*fliST* mutant was confirmed, and complementation of the double mutation restored the WT Succ^-^ phenotype (Fig S3I).

It is known that the IraP anti-adaptor controls succinate metabolism by modulating RpoS stability at the protein level [16]. The fact that we identified an *iraP*::Tn*5* mutant was an effective validation of the use of the *rpoS^2X^* genetic background for the transposon mutagenesis.

During the complementation experiments, we observed that the presence of chloramphenicol (Cm) mildly stimulated the growth of the WT strain (Fig S3J), an observation that will be investigated below.

Certain spontaneous mutations that stimulated growth with succinate did not reflect a typical loss-of-function scenario. For example, one of the spontaneous Succ^+^ mutants had an additional CTA codon resulting in an extra leucine between residues 239 and 240 of the transcriptional regulator OxyR (mutant *oxyR^mut^*) (Table 1). We also identified two Succ^+^ mutants with a single nucleotide polymorphism (SNP) located in the 5’-UTR of *dctA* (mutants *dctA^mut1^* and *dctA^mut2^*, Table 1), that encodes for the aerobic succinate transporter DctA [44]. Finally, two Succ^+^ mutants carried a SNP in the anti-Shine-Dalgarno sequence of the *rrsA* and *rrsH* genes that encode two 16S ribosomal RNAs; mutants *rrsA^mut^* and *rrsH^mut^* (Table 1). The function of the mutations associated with genes *oxyR*, *dctA*, *rrsA* and *rrsH* was confirmed by scarless genomic editing to generate exactly the same nucleotide changes in the WT background (Methods). All these reconstructed mutations caused the Succ^+^ phenotype (Fig 3).

In summary, we identified novel mutations that promote *Salmonella* growth upon succinate. These included mutations that involved the transcriptional regulators (RbsR, IscR, OxyR), RNA binding proteins (PNPase, Hfq and CspC), flagellar protein chaperones (FliS and FliT) and in ribosomal RNAs (RrsA and RrsH) (Fig 3). The eleven novel Succ^+^ mutations also promoted *Salmonella* growth upon fumarate or malate, suggesting that the regulatory systems play a general role in the de-inhibition of C_4_-dicarboxylate utilisation (Supplementary Fig S1D).

### Hfq, PNPase and their cognate sRNAs maintain the inhibition of succinate utilisation

Our discovery that the inactivation of the RNA binding proteins Hfq and PNPase promoted *Salmonella* growth upon succinate (Fig 3) led us to investigate the phenotype in more Δ*hfq* and Δ*pnp* mutants displayed a lag time of ∼5 hours and ∼15 hours, respectively (Fig 4A), prompting experiments to investigate the role of small regulatory RNAs (sRNAs) in the inhibition of succinate utilisation.

**Figure 4.**
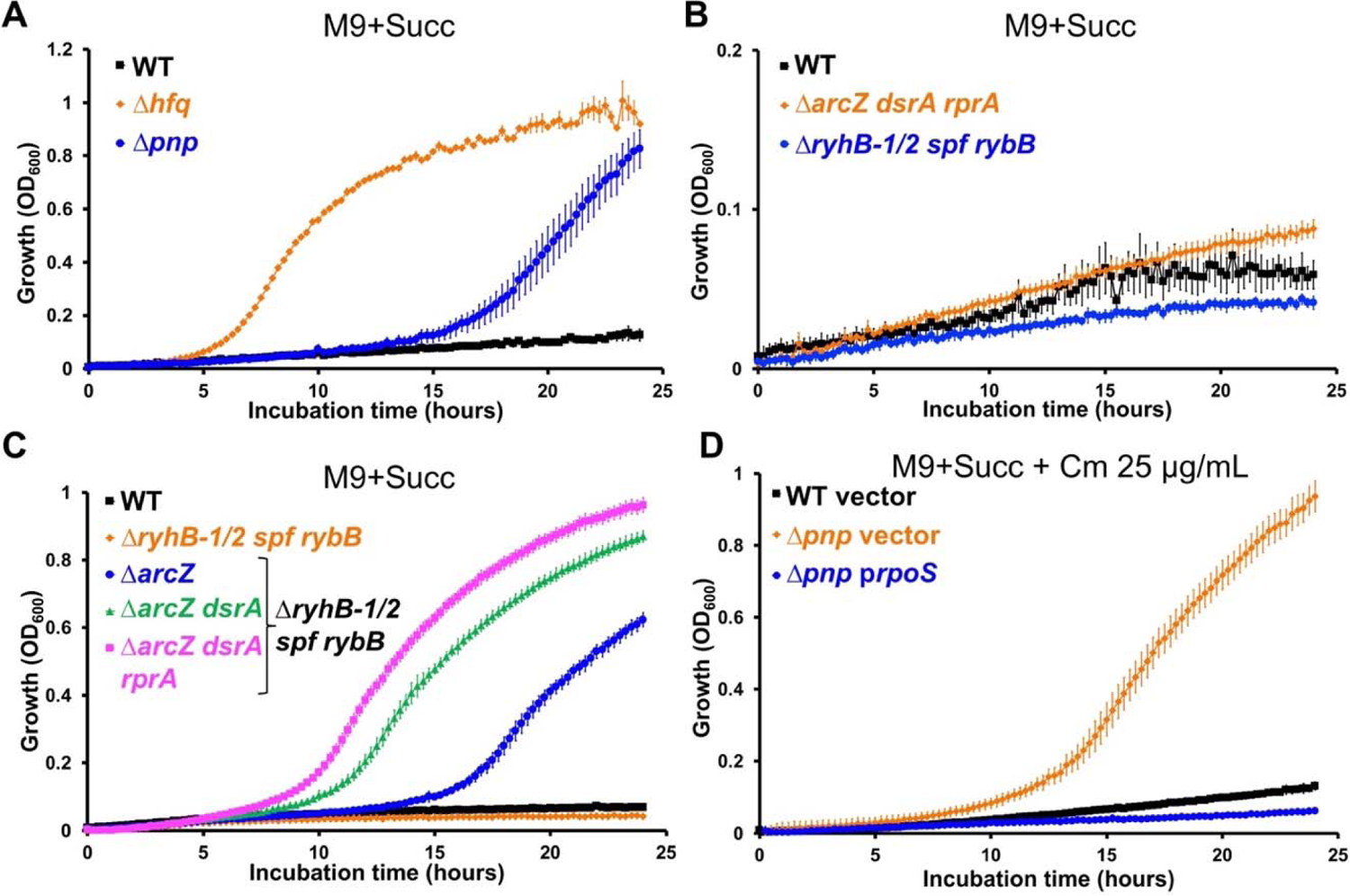
Hfq, PNPase and sRNAs inhibit *Salmonella* growth with succinate. (**A**) Hfq and PNPase inactivation boosts *Salmonella* growth on succinate. (**B**) The co-inactivations of the *rpoS* activating sRNAs ArcZ, DsrA and RprA or of the *sdh* repressing sRNAs RyhB-1/2, Spf and RybB did not stimulate *Salmonella* growth with succinate. (**C**) Successive inactivations of ArcZ, DsrA and RprA in the Δ*ryhB-1 ryhB-2 rybB spf* genetic background stimulate gradually the growth with succinate. (**D**) The overexpression of *rpoS* abolishes totally the Succ^+^ phenotype of the Δ*pnp* mutant, lacking PNPase: growth was assessed for strains 4/74 WT and Δ*pnp*, carrying the empty plasmid (vector, pNAW125) or the p*rpoS* (pNAW95) plasmid, overexpressing *rpoS*. The strains used were all 4/74 derivatives: Δ*hfq* (JH3584), Δ*pnp* (JH3649), Δ*arcZ dsrA rprA* (JH4385), Δ*ryhB-1 ryhB-2 rybB spf* (SNW630), Δ*ryhB-1 ryhB-2 rybB spf arcZ* (SNW639), Δ*ryhB-1 ryhB-2 rybB spf arcZ dsrA* (SNW640) and Δ*ryhB-1 ryhB-2 rybB spf arcZ dsrA rprA* (SNW641). The medium used is indicated for each experiment. Growth curves were carried out with 6 replicates grown in 96-well plates, as specified in Methods.

The RNA chaperone Hfq and its associated sRNAs are key post-transcriptional regulatory determinants [45, 46]. In *E. coli*, the sRNAs RybB, RyhB (RyhB-1 in *Salmonella*) and Spot42 (Spf) base-pair with the *sdhC* 5’-UTR to repress *sdhC* translation in an Hfq-dependent manner. In addition, RybB and RyhB reduce the stability of the *sdh* mRNA [47, 48]. We reasoned that the observed Succ^+^ phenotype of the Hfq null mutant could reflect de-repression of the *sdh* mRNA at the translational level.

In *E. coli*, the iron-dependent sRNA RyhB represses growth with succinate under iron-limited conditions [49]. Because exogenous iron was not added to our M9 media, we investigated whether the inhibition of *Salmonella* growth with succinate was the consequence of the *sdh* repression by RyhB-1 or RyhB-2, the RyhB-1 paralog in *Salmonella* [50]. Neither iron (FeCl_3_) supplementation (up to 100 µM) or the double inactivation of RyhB-1 and RyhB-2 (strain Δ *hB-1/2*) generated a Succ (Supplementary Fig S4). Similarly, the simultaneous inactivation of four sRNAs (RybB, Spf, RyhB-1 and RyhB-2) did not affect growth on M9+Succ (Fig 4B).

Hfq and sRNAs are crucial for the stimulation of *rpoS* translation. The long 5’-UTR of *rpoS* mRNA forms a self-inhibitory hairpin secondary structure, that blocks the ribosome access to the ribosome binding site, repressing *rpoS* mRNA translational initiation [51]. In *E. coli*, base-pairing of the sRNAs ArcZ, DsrA and RprA with the *rpoS* 5’-UTR, relieves this self-repression in an Hfq-dependent manner to stimulate *rpoS* translation [52]. In addition, binding of ArcZ, DsrA and RprA to the *rpoS* 5’-UTR prevents the premature Rho-dependent transcription termination of the *rpoS* mRNA [53]. As RpoS plays a pivotal role in the control of succinate utilisation, we assessed the growth of the triple Δ *Z rprA dsrA* mutant in M9+Succ. In comparison with the WT strain, no obvious differences were observed (Fig 4B). However, the successive deletion of sRNAs *arcZ*, *dsrA* and *rprA* in the Δ*rybB spf ryhB-1/2* genetic background did promote growth on succinate, and gradually reduced the duration of lag time (Fig 4C).

Inactivation of *pnp* is known to restore growth of a RyhB-overexpressing *E. coli* strain on succinate by reducing the stability of several sRNAs, including RyhB [54]. The same study demonstrated that the translational activation of *rpoS* by RprA and DsrA was attenuated in the Δ background. To test whether PNPase inactivation boosted succinate utilisation through RpoS attenuation, we assessed the growth of a Δ*pnp* mutant that overexpressed *rpoS*. The plasmid-borne overexpression of *rpoS* in this strain totally suppressed the Succ^+^ phenotype (Fig 4D), consistent with the stimulation of RpoS expression by PNPase.

Taken together, these results indicate that the fast growth of the Δ *q* and Δ*pnp* strains reflected both the dysregulation of the sRNA-mediated repression of *sdh* and the activation of *rpoS* translation. However, none of the sRNA mutants tested displayed the same fast-growing pattern of the Δ*hfq* mutant, suggesting that other sRNAs may be involved in the inhibition of succinate utilisation.

### The OxyS sRNA stimulates growth upon succinate by repressing expression of CspC

The spontaneous Succ^+^ mutants included an *oxyR* variant (*oxyR^mut^*) that encoded an extra leucine residue in the C-terminus domain of the OxyR transcriptional regulator (Table 1, Fig 5A). OxyR senses oxidative stress and is activated by disulfide bond formation in the presence of reactive oxygen species [55, 56]. In *E. coli*, the OxyR regulon includes about 40 genes, mainly associated with oxidative stress resistance [57, 58]. In addition, the oxidized form of OxyR triggers the transcription of OxyS, an Hfq-binding sRNA [59–61]. Previous studies reported the isolation of constitutively-active OxyR variants that carried mutations in the same region as the extra leucine residue of the *oxyR^mut^* variant [62–64]. Consequently, we investigated whether the *oxyR^mut^* allele drove constitutive transcription of the OxyS sRNA. Northern blot analysis revealed that OxyS was strongly expressed both in the absence and in the presence of hydrogen peroxide in the *oxyR^mut^* strain, indicating that the OxyR^mut^ protein is constitutively active (Fig 5B). We then determined whether OxyS constitutive expression was responsible for the Succ^+^ phenotype of the *oxyR^mut^* mutant. The deletion of *oxyS* in the *oxyR^mut^* strain (*oxyR^mut^* Δ*oxyS*) totally abolished the Succ^+^ phenotype (Fig 5C). A complementation experiment was carried out by re-introducing a single copy of *oxyS* and its native promoter (*oxyS^chr+^,* Fig S5) into the chromosome of the *oxyR^mut^* Δ*oxyS* strain (Methods). This chromosomal complementation restored the fast growth of the *oxyR^mut^* Δ*oxyS* mutant in M9+Succ (Fig 5C). Furthermore, the plasmid-borne expression of OxyS boosted the growth of 4/74 WT in M9+Succ, confirming that high level expression of the OxyS sRNA stimulated growth with succinate (Fig 5D). The same plasmid did not stimulate the growth of the Δ*oxyR* strain indicating that a functional OxyR is required for growth in M9+Succ (Fig 5D). We previously showed that Hfq inactivation boosted succinate utilisation (Fig 4A), but in the *oxyR^mut^* genetic background the same Hfq inactivation dramatically reduced growth and extended the duration of lag time in M9+Succ (Fig 5 E). Collectively, our findings show that the OxyS sRNA orchestrates the de-inhibition of succinate utilisation in an Hfq-dependent manner.

**Figure 5.**
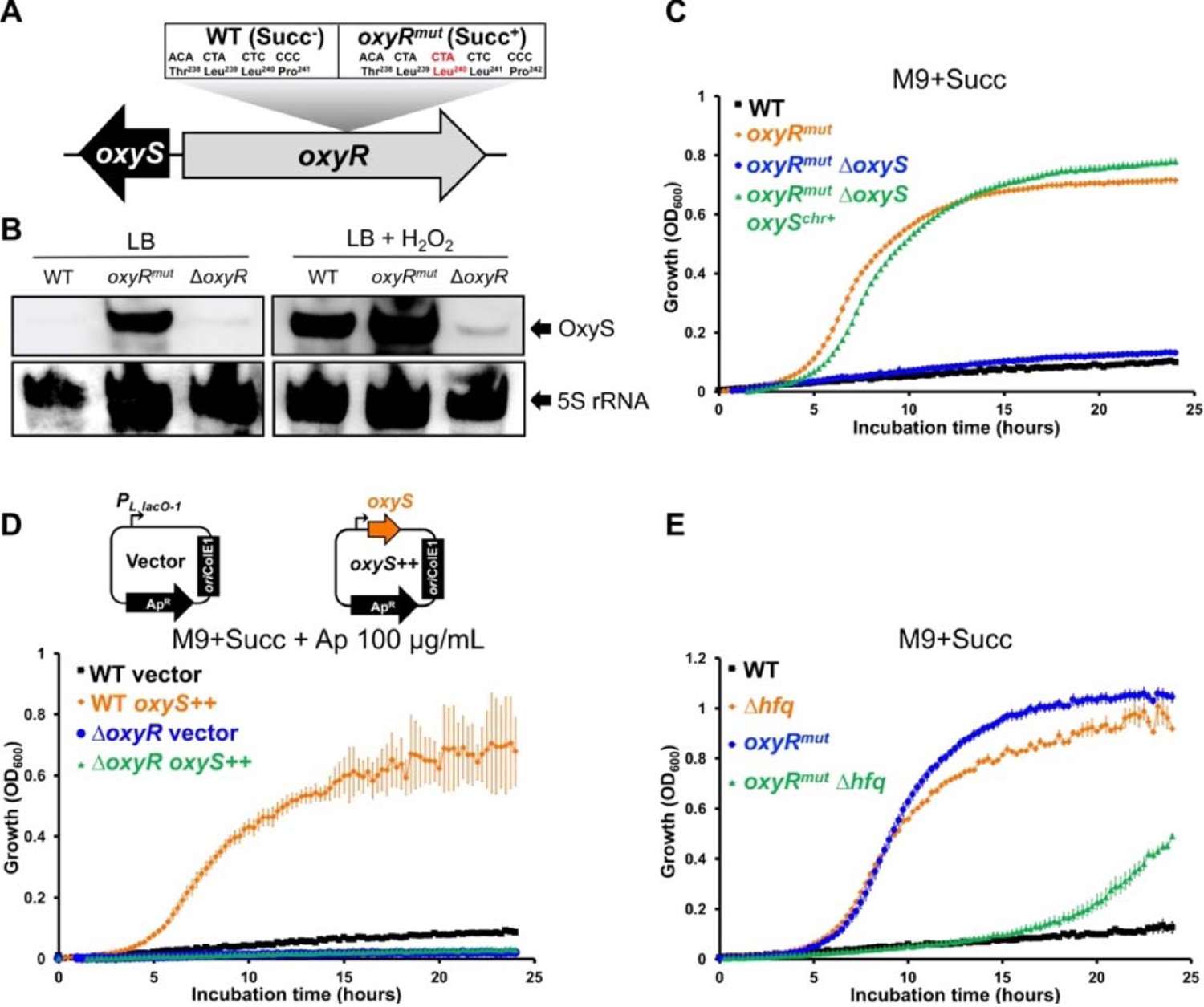
The Succ^+^ *oxyR^mut^* mutant expresses OxyS constitutively and stimulates *Salmonella* growth with succinate in an Hfq-dependent manner. (**A**) Schematic representation of the *oxyRS* locus. The Succ^+^ mutation *oxyR^mut^* has an additional CTA codon encoding for an extra leucine in *oxyR*. (**B**) OxyS is constitutively expressed in the *oxyR^mut^* strain. Northern blot detection revealed OxyS expression in strains 4/74 WT, *oxyR^mut^* (SNW318) and *oxyR* (SNW320) grown in LB and exposed or not to 2 mM H_2_O_2_ for 30 min (Methods). The detection of the 5S rRNA was used as a loading control. (**C**) OxyS stimulates *Salmonella* growth with succinate. Growth curves showed the fast growth of the *oxyR^mut^* strains in comparison with the WT and the *oxyR^mut^* Δ*oxyS* (SNW340) strains. Growth is restored in the complemented strain *oxyR^mut^* Δ*oxyS oxyS^chr+^* (SNW362), carrying a chromosomal *oxyS* copy. (**D**) The plasmid-borne expression of OxyS stimulates the growth of the WT but not of the Δ*oxyR* mutant. Growth was assessed for strains 4/74 WT and Δ*oxyR* (SNW320), carrying either the empty (vector, pP_L_) or the *oxyS* expressing (*oxyS++,* pNAW255) plasmids, schematised at the top of the figure. The bent arrows represent the constitutive promoter *P_L lacO-1_* of the Ap^R^ pP_L_ plasmid, carrying the *ori*ColE1 replicon. (**E**) Hfq inactivation suppresses the fast growth of the *oxyR^mut^* mutants. Growth was assessed for strains 4/74 WT, Δ*hfq* (JH3584), *oxyR^mut^* (SNW318) and *oxyR^mut^* Δ*hfq* (SNW663). The medium used is indicated for each experiment. Growth curves were performed in 96-well plates, as specified in Methods. The details about the construction of strains SNW320, SNW340 and SNW362 are depicted in the supplementary Fig S5.

In *E. coli*, OxyS acts as an indirect repressor of RpoS expression, probably *via* the titration of Hfq [60]. OxyS also represses the expression of the *yobF-cspC* operon, probably by base-pairing near the SD motif of the *yobF* 5’-UTR [65, 66]. Because RpoS, Hfq and CspC repress succinate utilisation (Fig 3), we tested the effects of the plasmid-borne overexpression of *rpoS^+^*, *hfq^+^* or *cspC^+^* on the growth of the *oxyR^mut^* strain. The overexpression of Hfq and RpoS slightly increased the lag time of *OxyR^mut^* strain, while the plasmid-borne expression of CspC totally abolished the Succ^+^ phenotype in this genetic background (Fig 6A).

**Figure 6.**
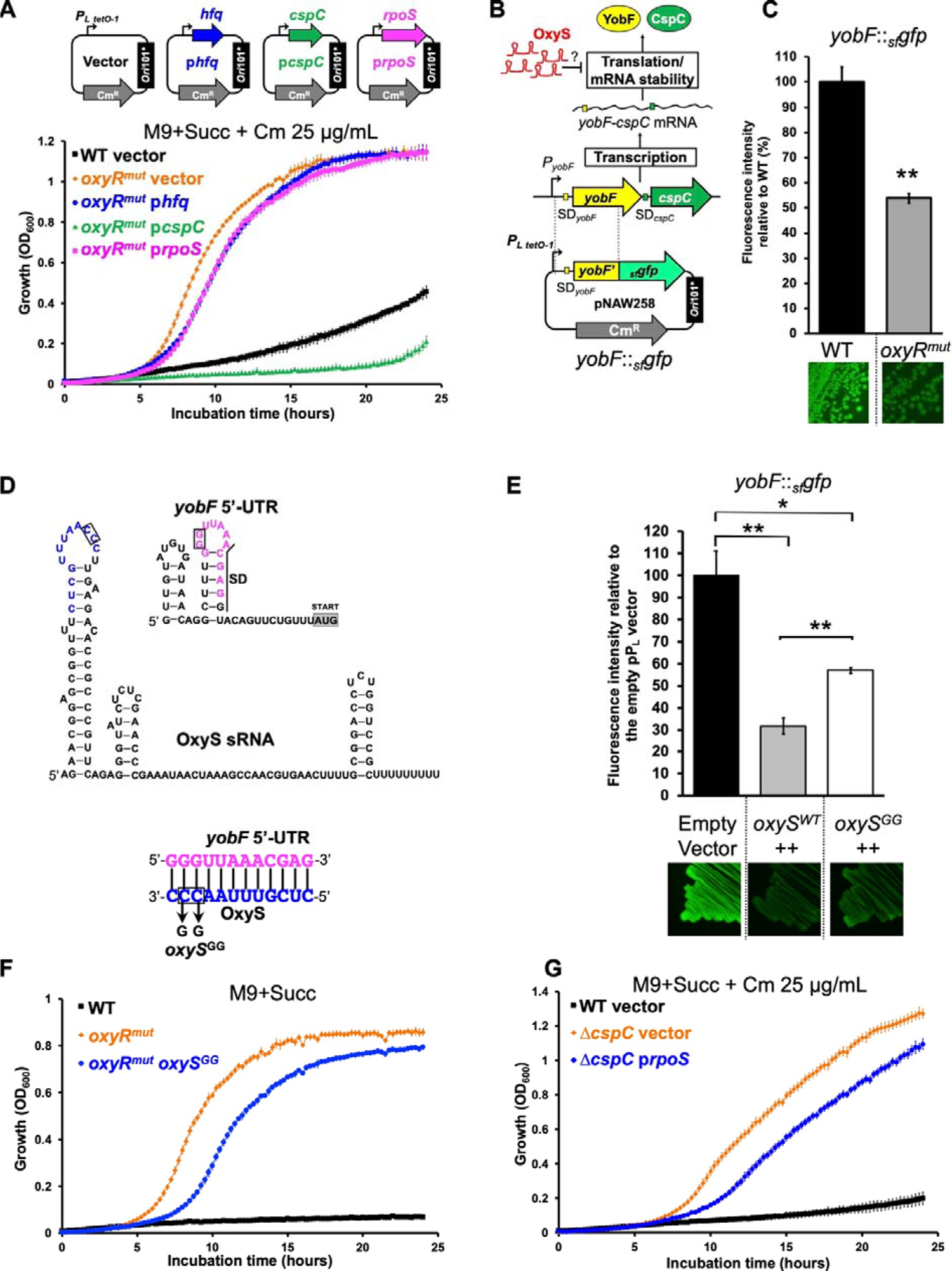
The OxyS sRNA stimulates *Salmonella* growth with succinate by repressing expression of CspC. (**A**) Overexpression of *cspC* inhibits the growth of the *oxyR^mut^* strain (SNW318), while *hfq* and *rpoS* overexpression have only a mild effect. The *oxyR^mut^* strain was carrying either the empty (vector, pNAW125), the p*rpoS* (pNAW95), the p*hfq* (pNAW45) or the p*cspC* (pNAW92) plasmids, schematised at the top of the figure. These low-copy Cm^R^ plasmids are carrying the *ori*101* replicon and the genes of interest are under the control of the strong constitutive promoter *P_L tetO-1_* (bent arrows). As a control, the growth of 4/74 WT carrying the empty plasmid (vector) was also assessed. (**B**) Strategy to test whether the *Salmonella* sRNA OxyS represses the expression of the *yobF-cspC* mRNA at the post-transcriptional level, as previously reported in *E. coli* [65]. The plasmid-borne translational fusion *yobF*::*_sf_gfp* (pNAW258) is depicted and was constructed as described in Methods and in Corcoran *et al.*, 2012 [121]. This fusion is under the control of the constitutive promoter *P_L tetO-1_*.SD= Shine-Dalgarno. (**C**) The *yobF*::*_sf_gfp* activity is reduced in the *oxyR^mut^* strain, that expresses constitutively OxyS: fluorescence GFP intensities were measured in strains 4/74 WT and *oxyR^mut^* carrying *yobF*::*_sf_gfp*. (**D**) Prediction of the RNA secondary structure of *Salmonella* OxyS and of *yobF* 5’-UTR, using Mfold [141]. The putative kissing complex between the two RNA molecules was predicted with IntaRNA [142] and the corresponding nucleotides are indicated in magenta for *yobF* 5’-UTR or in blue for OxyS. The mutation *oxyS^GG^* is indicated and the corresponding nucleotides are framed. (**E**) The plasmid-borne overexpression of OxyS represses the *yobF*::*_sf_gfp* activity and the *oxyS^GG^* mutation attenuates this repression: fluorescence GFP intensities were measured in the Δ *yS* strain (SNW338) carrying *yobF*::*_sf_gfp* and the empty (empty vector, pP_L_), the pP_L_-*oxyS* (*ox*yS*^WT^*++, pNAW255) or the pP_L_-*oxyS^GG^* (*ox*yS*^GG^*++, pNAW259) plasmids. (**F**) The *oxyS^GG^* mutation reduces the growth of the *oxyR^mut^* strain: strains 4/74 WT, *oxyR^mut^* and the *oxyR^mut^* mutant carrying the *oxyS^GG^* mutation (SNW670) were grown in M9+Succ. (**G**) Effects of *rpoS* overexpression on the growth of the Δ*cspC* (SNW292), carrying either the empty plasmid (vector, pNAW125) or the p*rpoS* (pNAW95) plasmid were grown in M9+Succ. For A, F and G, growth curves were carried out in the indicated medium with 6 replicates grown in 96-well plates. For C & E, strains were grown to OD_600_ ∼ 2 in LB, supplemented with the appropriate antibiotic(s). GFP fluorescence intensities were measured, as specified in Methods. The graphs represent the relative fluorescence intensities (%), in comparison with the indicated reference strain (100% of intensity). The same strains carrying GFP fusions were grown on LB agar plates and pictures were taken under blue light exposure. The data are presented as the average of biological triplicates ± standard deviation and the statistical significance is indicated, as specified in Methods.

To confirm that the OxyS-driven repression of the *yobF-cspC* was conserved in *Salmonella* we used a plasmid-borne *yobF*::*_sf_gfp* translational reporter (Fig 6B). In comparison with the WT, the *yobF*::*_sf_gfp* activity was significantly lower in the *oxyR^mut^* strain (∼2-fold repression), confirming that OxyS represses the expression of the *yobF-cspC* operon in *Salmonella* (Fig 6C). Bioinformatic analyses identified the putative secondary structures and the potential base-pairing interaction between OxyS and the *yobF* 5’-UTR (Fig 6D), which was predicted to be an 11 nucleotide-long kissing complex between OxyS and the *yobF* 5’-UTR, consistent with the proposed interaction in *E. coli* [66] (Fig 6D).

To assess the role of the kissing complex experimentally, we generated a mutated version of OxyS with a CC->GG mutation in the loop of the first RNA hairpin (allele *oxyS^GG^,* Fig 6D). This mutation was introduced into the chromosome of the *oxyR^mut^* strain (strain *oxyR^mut^ oxyS^GG^*) and the *oxyS^GG^* gene was cloned into the pP_L_ expression vector [67]. The empty pP_L_ vector, the pP_L_-*oxyS* or the pP_L_-*OxyS^GG^* plasmids were transferred into the Δ*oxyS* mutant carrying the *yobF*::*_sf_gfp* fusion and the GFP signal was measured. In comparison with the empty pP_L_ vector, in the presence of the pP_L_-*oxyS* (*oxyS++*) reduced the GFP fluorescence intensity by ∼3-fold, but only by ∼1.5-fold in the presence of the pP_L_-*oxyS^GG^* (*oxyS^GG^++*) (Fig 6E). Consistent with the attenuated repression of *yobF* in the presence of OxyS^GG^, the *oxyR^mut^ oxyS^GG^* strain had a longer lag time than the *oxyR^mut^* mutant, confirming that the mutated region of OxyS is involved in the de-inhibition of succinate utilisation (Fig 6F).

In *E. coli*, CspC stabilises *rpoS* mRNA and increases the cellular level of RpoS [68, 69]. To investigate whether the Succ^+^ phenotype of the CspC null mutant was caused by changes in RpoS expression, we tested the effect of RpoS overexpression in the Δ *pC* mutant (Fig 6G). The plasmid-encoded overexpression of RpoS only marginally extended lag time in Δ*cspC* mutant, indicating that repression of succinate utilisation by CspC is RpoS-independent. A recent study corroborated this observation, as the CspC-mediated activation of RpoS was not observed in *Salmonella* [70].

Collectively, our results indicate that the OxyS sRNA is a key determinant in the de-inhibition of succinate utilisation by *Salmonella*. Despite the fact that OxyS can regulate RpoS expression levels, we propose that OxyS stimulates the Succ^+^ phenotype by repressing the expression of CspC *via* base-pairing in the vicinity of the *yobF* SD motif.

CspC is an RNA binding protein, belonging to the cold shock protein family [71]. CspC and its paralog CspE often have redundant functions, being involved in biofilm formation, motility, stress resistance and virulence modulation in *S.* Typhimurium [68, 70]. It remains unclear how the OxyS-driven inhibition of CspC expression impacts upon the catabolism of succinate. One possibility is that CspC directly represses succinate catabolic genes. In line with this hypothesis, a transcriptomic study in *S.* Typhimurium, revealed that the *sdhC*, *sdhD* and *sdhA* genes are moderately up-regulated in a Δ*cspEC* mutant [70].

### The iron-sulphur cluster regulator IscR inhibits growth upon succinate by repressing DctA expression

In *E. coli*, the C_4_-dicarboxylate transporter DctA mediates succinate uptake under aerobic conditions [44]. DctA is also a C_4_-dicarboxylate co-sensor and modulates the expression of several genes, including *dctA* itself, in concert with the two-component system DcuR/S [72]. The transcription of *dctA* is controlled by catabolic repression and putative CRP binding sites, conserved in *Salmonella*, have been identified in the *dctA* promoter region [73, 74] (Fig 7A).

**Figure 7.**
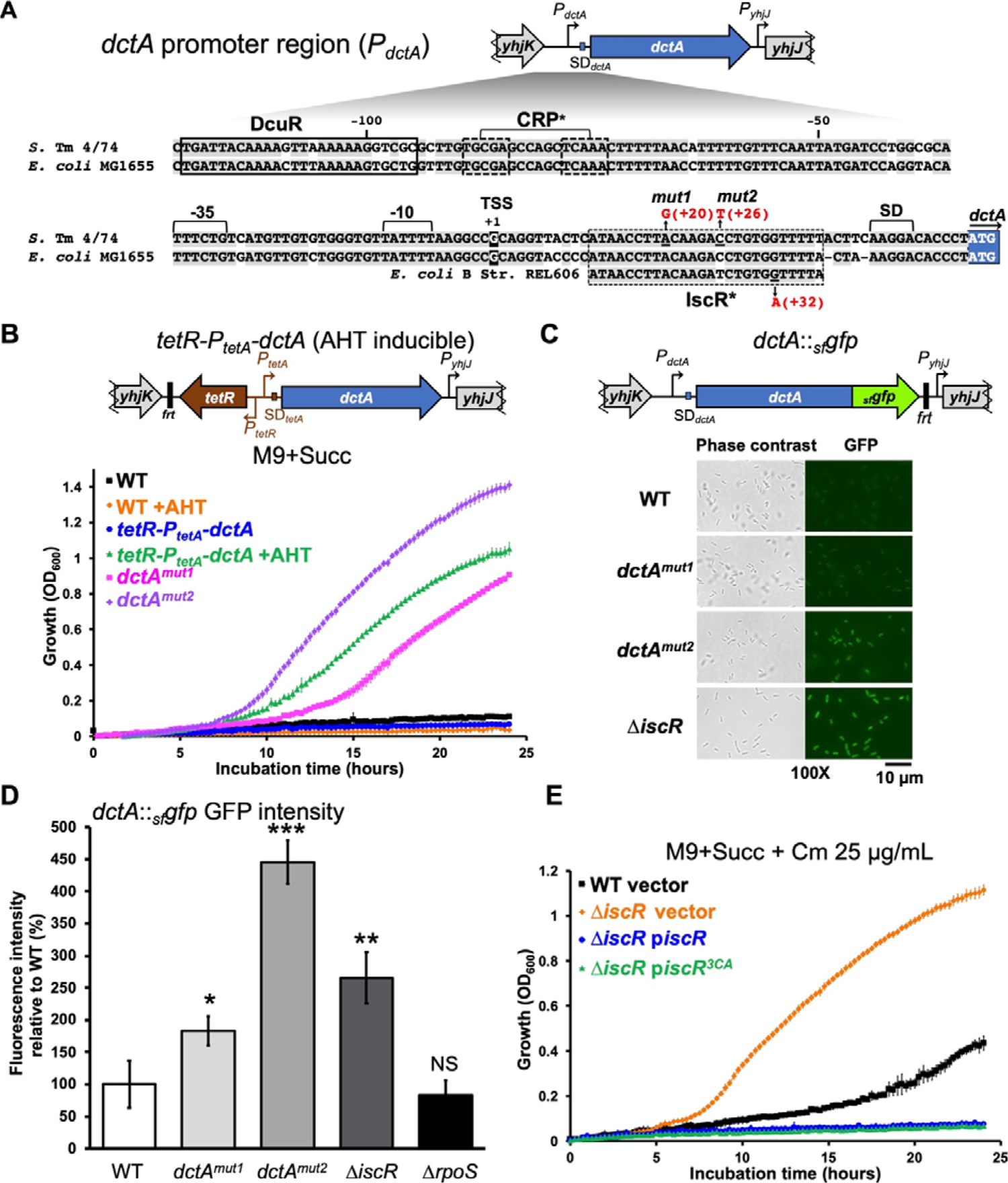
IscR represses the expression of DctA and inhibits *Salmonella* growth upon succinate. (**A**) Detailed schematic representation of the *dctA* promoter (*P_dctA_*) region in *S.* Typhimurium (*S.* Tm) 4/74 and in *E. coli* MG1655. Conserved nucleotides in 4/74 and MG1655 are highlighted in light gray. The promoter −35 and −10 boxes and the transcription start site (TSS, numbering +1) are indicated according to Davies *et al*., 1999 [73]. The DcuR binding site [143] and the putative CRP binding site [73, 74]. A putative IscR binding site is depicted and the mutations identified in the Succ^+^ mutants *dctA^mut1^* and *dctA^mut2^* are indicated in red. In addition, the corresponding region of *E. coli* B strain REL606 is depicted: in this strain, a G A SNP (in red) causing *dctA* up-regulation and the stimulation of succinate utilisation was previously described [81]. SD = Shine-Dalgarno motif. Promoters (*P*) are represented by bent arrows. The symbol “*” denotes that the binding sites were not experimentally demonstrated. (**B**) The stimulation of DctA expression and the *dctA^mut1^* and *dctA^mut2^* mutations boost *Salmonella* growth with succinate. The AHT-inducible strain *tetR-P_tetA_-dctA* (SNW133) is depicted: the *tetR* repressor gene, the *P_tetR_* and the *P_tetA_* promoters are indicated. The residual FLP recognition target site sequence is denoted by “*frt*”. In the absence of AHT no growth was detected for both WT and *tetR-P_tetA_-dctA* strains, while AHT addition stimulated the growth of the *tetR-P_tetA_-dctA* strain. Similarly, the *dctA^mut1^* (SNW160) and *dctA^mut2^* (SNW315) strains displayed a fast growth in M9+Succ. (**C** & **D**) The SNP mutations *dctA^mut1^* and *dctA^mut2^* and the inactivation of IscR stimulate the expression of DctA. The chromosomal transcriptional/translation *dctA*::*_sf_gfp* fusion is depicted: *_sf_gfp* encodes for the superfolder GFP fused in frame to DctA C-term. The GFP fluorescence intensity was measured in strain WT *dctA*::*_sf_gfp* (SNW296), *dctA^mut1^ dctA*::*_sf_gfp* (SNW310), *dctA^mut2^ dctA*::*_sf_gfp* (SNW316), Δ*iscR dctA*::*_sf_gfp* (SNW329) and Δ*rpoS dctA*::*_sf_gfp* (SNW313).(**B**) Both Apo- and holo-forms of IscR repress *Salmonella* growth with succinate. The growth was Δ*iscR* (SNW184) carrying either the empty plasmid (vector, pXG10-SF), the p*iscR* (pNAW96) or the p*iscR^3CA^* (pNAW97) plasmids. The p*iscR^3CA^* expressed the IscR^3CA^ variant that prevents the binding of an iron-sulphur cluster, maintaining IscR in its apo-form (see text for details). Growth curves (**B** & **E**) were carried out in the indicated medium with 6 replicates grown in 96-well plates. For **C** & **D**, strains carrying the *dctA*::*_sf_gfp* fusion were grown in M9+Gly+Succ to OD_600_ 0.5-1 and GFP fluorescence intensity was measured by fluorescence microscopy (**C**) or by flow cytometry (**D**), as specified in Methods. The graph (**D**) represents the relative fluorescence intensities of each strain (%), in comparison with the WT strain carrying *dctA*::*_sf_gfp* (SNW296, 100% of intensity). The data are presented as the average of biological triplicates ± standard deviation and the statistical significance is indicated, as specified in Methods. NS, not significant.

To determine whether DctA was required for *Salmonella* growth under our experimental conditions, we constructed a chromosomal inducible *dctA* construct, by replacing the *dctA* promoter with *tetR* (encoding the TetR repressor) and the *tetA* promoter (strain *tetR-P_tetA_-dctA,* Fig 7B). In the absence of anhydrotetracycline (AHT) inducer, the *tetR-P_tetA_-dctA* strain did not grow at all in M9+Succ. However, upon addition of AHT, the *tetR-P_tetA_-dctA* strain displayed a fast growth phenotype that was not observed with the WT (Fig 7B). We conclude that *Salmonella* requires the DctA transporter to grow on succinate and that *dctA* expression is likely to be repressed in the WT, as previously hypothesised by Hersch and co-workers [16].

The two spontaneous Succ^+^ mutants *dctA^mut1^* and *dctA^mut2^* (Table1) carry SNPs in the 5’-UTR of *dctA* (Fig 7A), and both promoted growth with succinate (Fig 7B). We reasoned that the Succ^+^ mutations could de-repress *dctA* expression, which we examined with a chromosomal *dctA*::*_sf_gfp* transcriptional/translational reporter fusion in the Succ^+^ mutant backgrounds (Methods and Fig 7C). To allow homogenous growth for all the strains (including the Succ^-^ 4/74 WT), bacteria were grown in M9 supplemented with both glycerol (40 mM) as the main C-source and the addition of 10 mM succinate, to stimulate expression of succinate-induced genes [75]. The GFP fluorescence intensity of single bacteria was measured for each strain by flow cytometry. In comparison with low levels of GFP fluorescence seen in the 4/74 WT background, higher GFP levels were only observed in the presence of the *dctA^mut1^, dctA^mut2^* and Δ*iscR* mutations (Supplementary Fig S6 A-N). The regulation was confirmed by fluorescence microscopy (Fig 7C) and flow cytometry, using biological triplicates (Fig 7D).

A recent report proposed that RpoS indirectly represses *dctA* in *Salmonella* [16], but such an increase of the *dctA*::*_sf_gfp* fusion activity was not observed in the Δ *rpoS* strain under our experimental conditions (Fig 7D, Supplementary Fig S6 P). Of note, the *dctA*::*_sf_gfp-*tagged Δ *rpoS*, *dctA^mut1^, dctA^mut2^* and Δ*rpoS*, *dctAmut1, dctAmut2* and Δ*iscR* mutants grew much faster in M9+Succ than the isogenic WT strain (Supplementary Fig S6O), with the same growth rate we observed previously with the corresponding untagged mutants (Fig 7B&E and Fig S1B). This indicated that the C-terminal addition of sfGFP did not impede the function of DctA as a succinate transporter or co-sensor.

The up-regulation of *dctA* in the *Salmonella* IscR null mutant was consistent with the IscR-driven repression of *dctA* proposed in *E. coli* [76]. In most Gram-negative bacteria, the dual regulator IscR controls the transcription of the iron-sulphur (Fe-S) cluster biosynthesis operon *iscRSUA* and the sulphur mobilization genes *sufABCDSE* [77, 78]. The apoprotein form of IscR (apo-IscR) is matured by the Isc system into its [Fe_2_-S_2_]-containing holo-form. The resulting holo-IscR represses the expression of several genes including the *isc* operon. Under iron-limitation and in the presence of reactive oxygen species, the IscR apo-form predominates and stimulates the expression of the *suf* operon, in concert with OxyR [77, 79]. IscR binds to two classes of DNA motifs: the Type 1 motifs are only bound by holo-IscR, while the Type 2 motifs are recognised by both holo- and apo-forms [76, 80].

Analysis of the promoter region of *dctA* revealed the presence of a putative Type 2 IscR-binding site (ATAACCTTACAAGACCTGTGGTTTTT) [80] located 10 bp downstream of the transcription start site of *dctA* (Fig 7A). Both the Succ^+^ mutants *dctA^mut1^, dctA^mut2^* carried SNPs within this DNA motif. This motif is also conserved in *E. coli* MG1655 (Fig 7A) and a similar SNP, that stimulated *dctA* transcription and succinate utilisation, was previously identified in the *E. coli* B strain REL606 [81] (Figure 7A).

To investigate which of the apo/holo-forms of IscR was repressing succinate utilisation in *Salmonella*, we constructed a plasmid expressing an IscR variant carrying three Cys Ala substitutions (IscR^3CA^,Cys_92,98,104_ Ala_92,98,104_) that prevent the binding of [Fe_2_-S_2_] to IscR, and maintain the apo-form of the protein [82, 83]. The plasmid-borne expression of both IscR and IscR^3CA^ complemented the Δ*iscR* deletion and suppressed the Succ^+^ phenotype (Figure 7 E), indicating that both apo- and holo-IscR repress succinate utilisation.

Collectively, these results demonstrated that IscR plays a critical role in the repression of *dctA* and in the inhibition of succinate utilisation. The apo-IscR represses succinate utilisation, suggesting that IscR represses *dctA* expression by binding the putative Type 2 DNA motif identified downstream of the *dctA* promoter. The finding of *dctA-*stimulating SNPs in this motif supports our hypothesis, and it is possible that the binding of IscR downstream of the promoter acts as a “roadblock” that inhibits *dctA* transcription, as proposed for the repression of *mgtC* by PhoP [84]. However, we cannot rule out that the *dctA* repression by IscR is indirect with the *dctA^mut1^* and *dctA^mut2^* mutations stimulating *dctA* transcription by another mechanism. Further study is required to understand the IscR-driven repression of *dctA* at the mechanistic level.

Our data show that the sole de-repression of *dctA* expression mutations is sufficient to stimulate *Salmonella* growth with succinate, consistent with a previous report [16]. However, *dctA* de-repression was only observed in the *dctA^mut1^, dctA^mut2^* and Δ*iscR* mutants, raising the question of whether DctA-driven succinate uptake is the key limiting factor for rapid growth of *Salmonella* in M9+Succ or whether overexpression of the DctA transporter, in its role of succinate co-sensor, may indirectly boost the expression of other limiting succinate utilisation genes.

### Succinate utilisation is inhibited by RbsR and FliST *via* RpoS

Factors that modulate RpoS expression, stability or activity are likely to control succinate utilisation in *Salmonella*. For example, the inactivation of the anti-adapter IraP stimulates succinate utilisation by increasing RssB-facilitated proteolysis of RpoS by the ClpXP protease [16,85,86]. In our genetic screens we found that succinate utilisation was stimulated by the absence of IraP, CspC, RbsR and FliST and by increased expression of OxyS.

To assess RpoS levels in the corresponding Succ^+^ mutants, we used Western blot detection (Supplementary Fig S7A). The RpoS levels in the Δ*cspC*, Δ*fliST*, *oxyR^mut^* mutants were similar to the WT strain, while lower levels of RpoS were observed in the Δ*iraP* and Δ*rbsR* mutants. We confirmed the role of RbsR in RpoS activation by complementing the Δ*rbsR* mutation with the p*rbsR* plasmid, revealing that inactivation of RbsR reduced RpoS abundance in exponential and early stationary phases, but not in stationary phase (Fig 8A, Supplementary Fig S7B). During the characterisation of the Succ^+^ mutants, we noticed that the RbsR null strain was impaired in its capacity to form red, dry and rough colonies (RDAR), another RpoS-dependent phenotype of *Salmonella* [87]. The RDAR morphotype was restored in the Δ*rbsR* mutant by complementation with the plasmid p*rbsR* (Fig 8B). We observed that the plasmid-borne overexpression of RpoS in the RbsR null mutant totally abolished the Succ^+^ phenotype, indicating that RbsR represses succinate utilisation *via* the activation of RpoS (Fig 8C).

**Figure 8.**
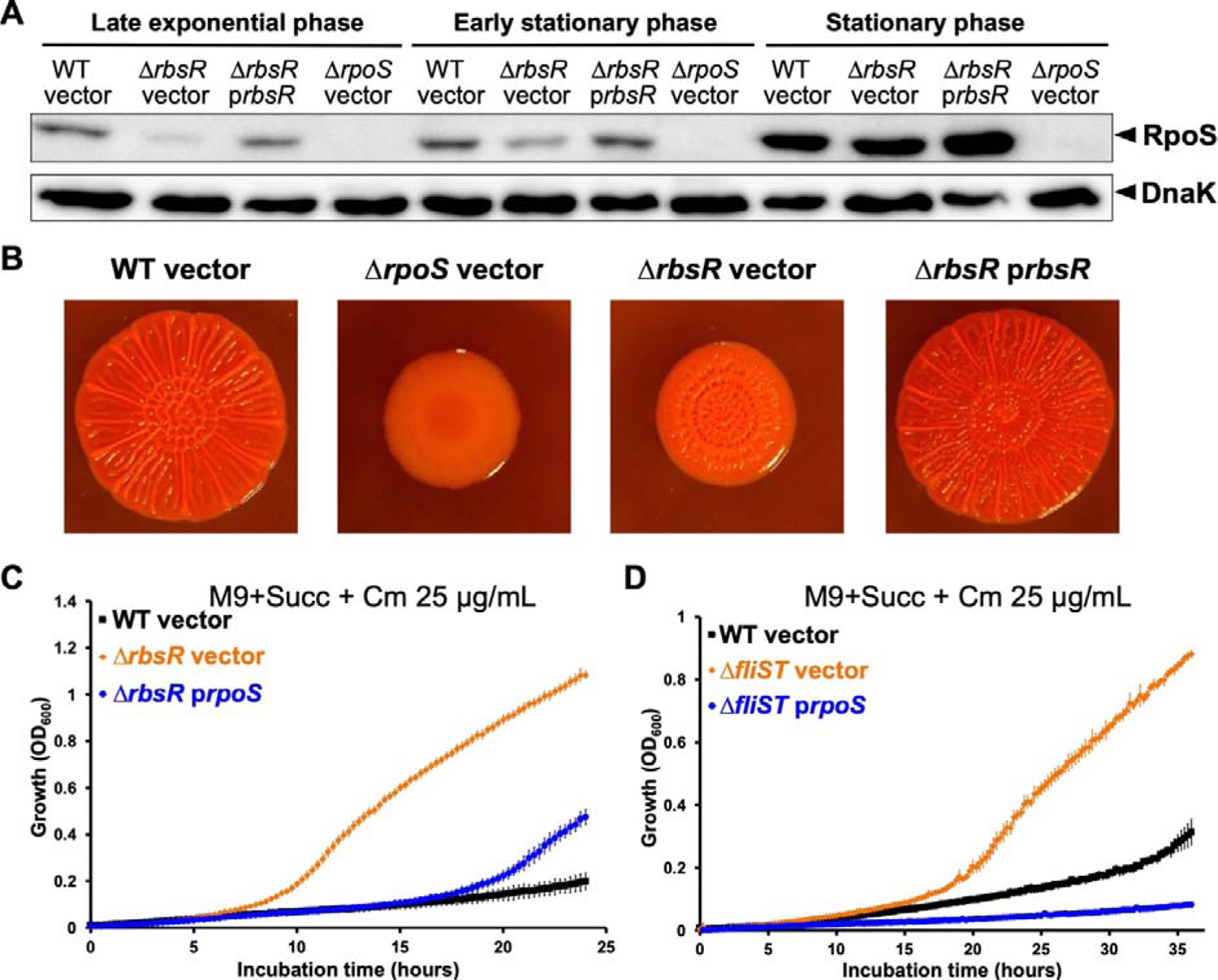
RbsR and FliST inhibit *Salmonella* growth upon succinate *via* RpoS. (**A**) RbsR inactivation reduces the cellular level of RpoS in exponential and early stationary phases. Strains 4/74 WT and Δ*rbsR* (SNW294) carrying the empty (vector, pXG10-SF) or the p*rbsR* (pNAW93) plasmids were grown in LB (without Cm) to late exponential (OD_600_∼1), early stationary (OD_600_∼2) and stationary phase (OD_600_∼4). The cellular levels of RpoS and DnaK (loading control) were assessed by Western blotting (Methods). As negative control, the Δ*rpoS* mutant (JH3674) carrying pXG10-SF was included. The experiment presented is representative of three independent experiments and two replicates are presented in Fig S7B. (**B**) RbsR inactivation reduces red, dry and rough (RDAR) morphotype, another RpoS-dependent phenotype. The RDAR phenotypic assays were carried out as specified in Methods with strain 4/74 WT, Δ*rbsR* and Δ*rpoS* carrying the indicated plasmids (vector = pXG10-SF). At least three independent experiments were performed, and representative RDAR colony pictures are presented. The plasmid-borne *rpoS* overexpression suppresses the Succ^+^ phenotype of the Δ*rbsR* (**C**) and Δ*fliST* (**D**) mutants. Growth was assessed in the indicated medium in a 96-well plate for strains Δ*rbsR* (SNW294) or Δ*fliST* (SNW288) carrying either the empty plasmid (vector, pNAW125) or the p*rpoS* plasmid (pNAW95) and strains 4/74 WT carrying the empty plasmid.

RbsR is a LacI-type transcriptional regulator that inhibits the transcription of the ribose utilisation operon (*rbsDACBKR*) in the absence of ribose [88]. To investigate whether RbsR stimulated *rpoS* transcription, we used a chromosomal transcriptional GFP reporter fusion, where the *gfp*^+^ gene (including its SD sequence) was inserted downstream of the main transcription start site of the *rpoS* locus (Fig S7C). Similar GFP levels were observed in the WT and the Δ*rbsR* strains grown to either exponential, early stationary or stationary phase (Fig S7D).

Taken together, our findings show that RbsR acts as a pleiotropic regulator in *Salmonella*, controlling growth with succinate and the RDAR morphotype, *via* the positive regulation of *rpoS*. RbsR does not directly stimulate *rpoS* promoter activity and we propose an indirect RbsR-driven activation of RpoS at the post-transcriptional or the post-translational level. In line with this hypothesis, it was recently observed that RpoS is repressed at the post-transcriptional level, when *rbsD*, a gene controlled by RbsR, is over-expressed in *E. coli* [89].

Our work also revealed that the two flagellar chaperones FliS [90] and FliT [91] control succinate utilisation (Fig 3, Fig S2B), suggesting a link between the control of the flagellar machinery and *Salmonella* central carbon metabolism. In the Δ*fliST* mutant, RpoS overexpression totally suppressed the Succ^+^ phenotype (Fig 8C) indicating that the regulation is RpoS-dependent. However, Western blots did not show reduced levels of RpoS in the FliST null mutant (Fig S7A). It remains unclear how these protein chaperones inhibit succinate utilisation, and whether FliS and FliT are capable of stimulating the expression or the activity of RpoS.

### Anti-Shine-Dalgarno mutations and sub-inhibitory concentration of chloramphenicol boost succinate utilisation

We identified a novel class of mutations that boost succinate utilisation by altering the anti-Shine-Dalgarno sequence (aSD) of the 16S ribosomal RNAs (rRNAs). Specifically, aSD SNPs in *rrsA* and *rrsH* genes that encode two of the seven 16S rRNAs present in *Salmonella* genomes [92] were found (alleles *rrsA^mut^* and *rrsH^mut^*, Fig 9A). Mature 16S rRNAs are assembled with ribosomal proteins to form the 30S ribosomal subunits that initiate mRNA translation [93, 94]. Each 16S rRNA 3’-end carries an aSD motif (CCUCCUU) that base-pairs with the Shine-Dalgarno sequence (SD) on mRNA, promoting translational initiation at the start codon [95, 96].

**Figure 9.**
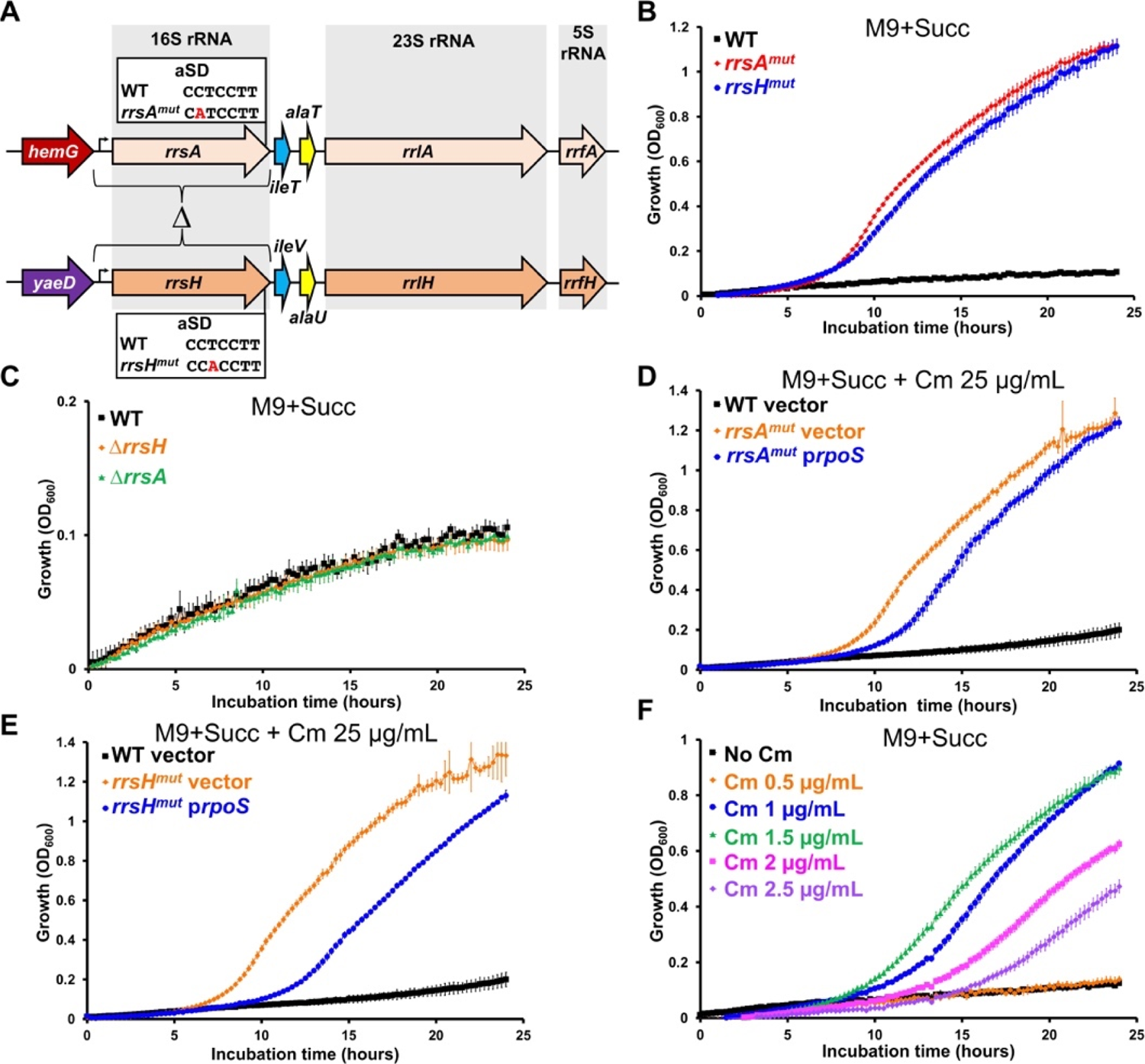
Mutation of the 16S rRNA aSD motifs and sub-inhibitory concentrations of chloramphenicol stimulate *Salmonella* growth upon succinate. (**A**) Schematic representation of the *Salmonella rrnA* and *rrnH* ribosomal RNA (rRNA) operons. The 23S (*rrl*), 16S (*rrs*) and 5S (*rrf*) rRNAs and the *ileT*, *ileV*, *alaT* and *alaU* tRNAs are represented, according to the annotation of the corresponding loci of *S.* Typhimurium LT2 (Genbank AE006468.2)[144]. The bent arrows represent the ribosomal promoter. The replacement of the full *rrsA* and *rrsH* loci (promoters included) with an I-*Sce*I-Km cassette (Methods) in strains Δ*rrsA* (SNW335) and Δ*rrsH* (SNW311) is represented by the “” symbol. The SNP mutations in the anti-shine-Dalgarno (aSD) motifs of mutant *rrsA^mut^* (SNW336) and *rrsH^mut^* (SNW314) are indicated in red. (**B**) The aSD mutations *rrsA^mut^* and *rrsH^mut^* stimulate *Salmonella* growth with succinate, while the full inactivation of the *rrsA* and *rrsH* loci (strains Δ*sA* and Δ*H*) did not affect the growth (**C**). The plasmid borne overexpression of *rpoS* has moderate effects on the growth of the *rrsA^mut^* (**D**) and *rrsA^mut^* (**E**) mutants with succinate. The 4/74 WT and the *rrs* mutants carried the empty plasmid (Vector, pNAW125) or the p*rpoS* (pNAW95) plasmid. (**F**) Subinhibitory concentration of chloramphenicol (Cm) stimulate *Salmonella* growth with succinate. All the growth curves were carried out with 6 replicates in 96-well plates with the indicated medium.

The SNPs carried by the *rrsA^mut^* and *rrsH^mut^* strains dramatically stimulated growth of *Salmonella* in M9+Succ, reducing the lag time to ∼7 hours. When *E. coli* grows under nutrient limitation, the relative transcription of the *rrnH* rRNA operon increases and the resulting pool of RrsH-containing ribosomes can modulate the stress response by stimulating RpoS translation or stability [97]. Therefore, we reasoned that the aSD mutations may totally inactivate the rRNAs resulting in the reduction of RpoS expression. However, deletion of the *rrsA* and *rrsH* loci did not result in a Succ^+^ phenotype (Fig 9C). The plasmid-borne expression of *rpoS* only marginally increased the lag time of the *rrsA^mut^* and *rrsH^mut^* mutant strains, indicating that the mutations in the 16S rRNAs stimulate succinate utilisation, at least partially, in a RpoS-independent manner (Fig 9D-E).

In *E. coli*, 16S rRNAs that carry a mutated aSD motif are processed and assembled into functional 30S subunits, which can initiate translation at the correct start codon [98]. This suggests that the mutated 16S rRNA RrsA^mut^ and RrsH^mut^ are assembled normally, and the presence of the resulting altered ribosome stimulates *Salmonella* growth upon succinate.

The aSD mutations prompted us to experiment with a translational inhibitor. We observed that subinhibitory concentrations of chloramphenicol (Cm) stimulated growth of 4/74 WT upon succinate (Fig 9F). The shortest lag time (∼8 hours) was observed at a Cm concentration of 1.5 µg/mL. Addition of Cm caused a similar level of growth stimulation for *S.* Enteritidis strain P125109 (Supplementary Fig S8A), indicating that the phenomenon is conserved in other *Salmonella* serovars.

Cm targets the 50S ribosome subunits to block translation [99]. Subinhibitory concentrations of this antibiotic prevent the RelA-mediated synthesis of the alarmone (p)ppGpp, the key signal molecule of the stringent response [100]. During amino acid starvation, (p)ppGpp accumulation is known to promote the transcription, translation and stability of RpoS [51], raising the possibility that the aSD mutations and Cm stimulate succinate utilisation directly through RpoS attenuation. Tetracycline (Tc) and other translation-inhibiting antibiotics also inhibit (p)ppGpp synthesis in *E. coli* [101], prompting us to test subinhibitory concentrations of Tc hydrochloride (1 and 2 µg/mL). However, Tc did not stimulate the growth of 4/74 at the concentrations tested (Fig S8B), suggesting that Cm does not reduce RpoS expression, *via* the inhibition of the stringent response.

Taken together, our findings suggest that the impairment of the ribosomal machinery by aSD mutations or by the presence of chloramphenicol impose a translational stress that stimulates genes involved in succinate utilisation. In line with this hypothesis, the inactivation of the translational elongation factor EF-P [102, 103] also stimulated the growth of *Salmonella* upon succinate [16]. However, the link between protein biosynthesis impairment and the stimulation of succinate utilisation remains enigmatic. Further work will be required to decipher the regulatory mechanism that underpins this phenomenon.

### Perspective

During infection, *Salmonella* and other pathogens face a metabolic dilemma between self-preservation and nutritional competence that is exemplified by succinate utilisation [1,104,105]. The RpoS master regulator functions as a double-edged sword, activating critical resistance mechanisms required for survival in the host [19, 20], and reducing nutritional capacity by repressing the utilisation of several infection-relevant C-sources, including succinate [22, 25].

Our genetic dissection revealed that RbsR, PNPase, Hfq and sRNAs modulate succinate metabolic capacity, *via* the fine-tuned control of RpoS. Furthermore, succinate utilisation is inhibited by the RpoS-independent CspC and IscR systems. These distinct regulatory mechanisms are likely to adjust *Salmonella* metabolism during the journey of the pathogen through the host; from the colonisation of the gastrointestinal tract to intra-macrophage replication. We showed that the sRNA OxyS antagonises CspC-dependent inhibition (Fig 5 & Fig 6). Because OxyS is induced by oxidative stress, our findings raise the possibility that the reactive oxygen species produced in the inflamed gut [106, 107] stimulate *Salmonella* growth upon microbiota-derived succinate in this niche [3, 5].

Despite, the abundance of succinate within infected macrophages [11], the intracellular proliferation of *Salmonella* does not require succinate catabolic genes [14]. The high levels of intra-macrophage expression of the *iscR* [108] and *rpoS* [108, 109] lead us to propose that the DctA-driven uptake and catabolism of succinate are strongly repressed in this cellular niche. Because succinate triggers the induction of *Salmonella* genes associated with survival and virulence within macrophages [13], we speculate that succinate utilisation is comprehensively repressed to prevent depletion of this critical signalling molecule from the intracellular niche.

Pioneering work from the 1960s established that constitutive succinate utilisation ablated *Salmonella* virulence in the murine infection model [110, 111], giving the first suggestion that tight regulation of succinate utilisation was critical for pathogenesis. Six decades later we have revealed that multiple systems control the utilisation of succinate, making the catabolism of succinate responsive to various environmental stimuli (Fig 10). We propose that the redundant regulatory systems ensure that *Salmonella* only utilises succinate “at the right place and at the right time” during infection.

**Figure 10.**
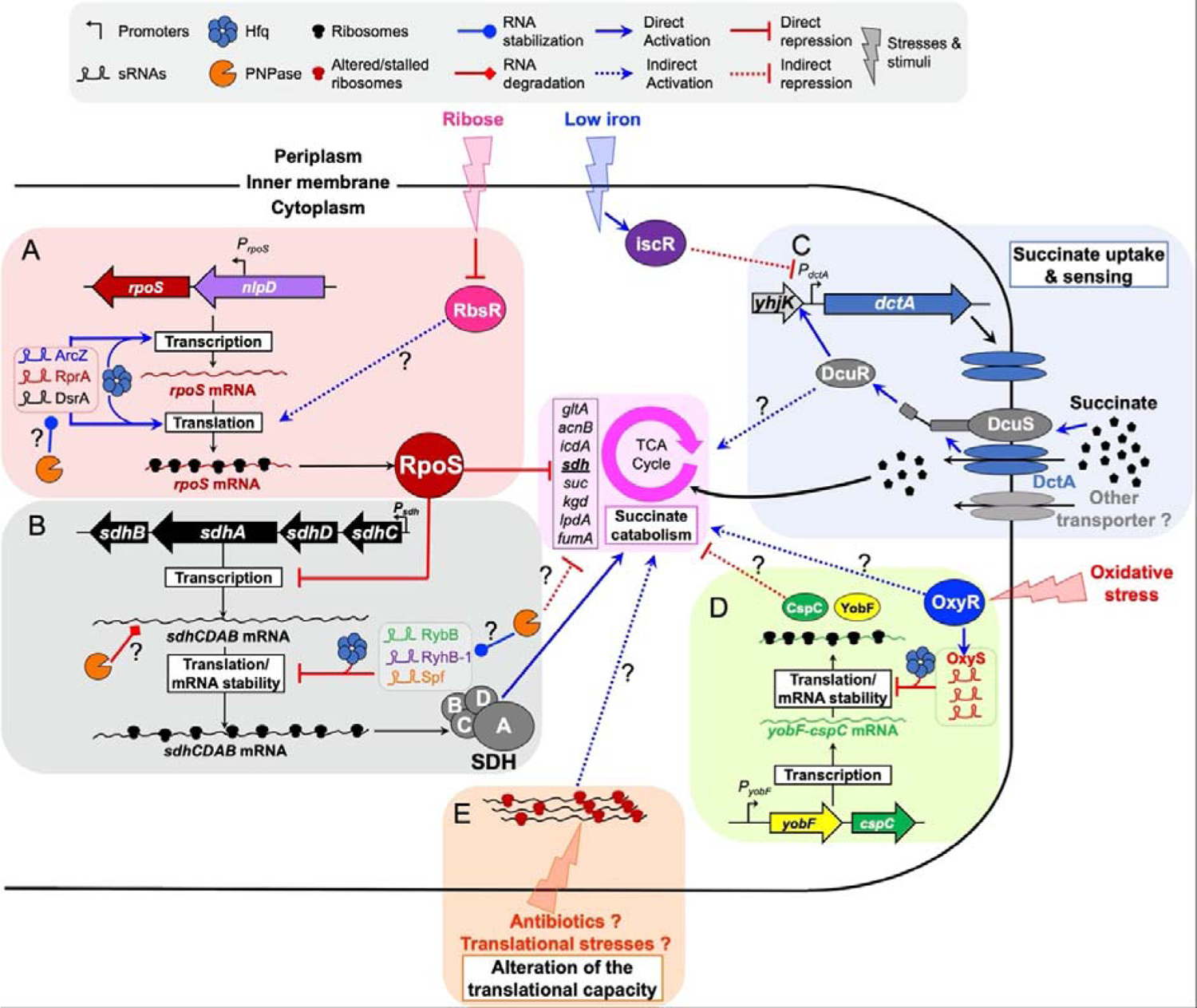
A model depicting the modulation of *Salmonella* succinate utilisation by multiple environmental stimuli. (**A**) RpoS expression stimulation by Hfq, PNPase, sRNAs and RbsR inhibit *Salmonella* succinate utilisation. The sRNAs ArcZ, DsrA and RprA stimulate *rpoS* mRNA transcription elongation [53] and translation [52] in concert with Hfq. RpoS represses the transcription of several genes of the TCA cycle, including the *sdh* operon [24, 25], inhibiting succinate catabolism and *Salmonella* growth with this C-source. PNPase presumably represses succinate utilisation indirectly by its role in the stabilization of several Hfq-associated sRNAs [145] and in the translational activation of *rpoS* [54]. The ribose sensor RbsR stimulates the expression of RpoS, presumably at the post-transcriptional level (Fig 8), repressing growth upon succinate. (**B**) The sRNAs RyhB-1, Spf and RybB repress *sdh* mRNA translation [47, 48] in concert with Hfq and attenuate succinate dehydrogenase (SDH) synthesis, inhibiting *Salmonella* growth upon succinate. In addition, PNPase may promote the degradation of the *sdhCD* mRNA [146]. (**C**) Under aerobic conditions, succinate is mainly imported by DctA [44]. DctA interacts with the DcuS protein and acts as a co-sensor of C_4_-dicarboxylates [72, 147]. In the presences of succinate, DcuS/DctA activates the response regulator DcuR, that stimulates the expression of several genes, including *dctA* [143]. *De novo* synthetised DctA accumulates and increases the uptake of succinate and DctA accumulation may also stimulate the transcription of succinate utilisation genes in concert with DcuS/R. However, in *Salmonella*, *dctA* expression is robustly repressed by the both halo- and apo-forms of the iron-sulphur cluster regulator IscR (Fig 7), which is up-regulated under iron limiting conditions [133]. Therefore, IscR plays a pivotal role in the succinate utilisation repression and blocks *Salmonella* growth with this C-source (**D**) OxyR is stimulated by oxidative stress and stimulates the expression of the sRNA OxyS. OxyS stimulates *Salmonella* succinate utilisation by repressing the small RNA binding protein CspC. CspC represses succinate utilisation by a still unknown mechanism. (**E**) Stressors that alter *Salmonella* translational capacity (*i.e.* 16S rRNAs mutations, antibiotics or environmental stressors) stimulate succinate utilisation by a still unknown mechanism. Interrogation marks indicate speculative interactions.

## Materials and Methods

### Bacterial strains and growth conditions

Precise details of all the chemicals, reagents, DNA oligonucleotides (primers), plasmids and bacterial strains used in this study are listed in Supplementary Resource Table S1. The *Salmonella* mutant strains were all derivatives of *Salmonella enterica* serovar Typhimurium strain 4/74 [112]. Strain 4/74 is now available from the UK National Collection of Type Cultures (https://www.culturecollections.org.uk/products/bacteria/index.jsp) as NCTC 14672. All the nucleotide coordinates given for 4/74-derived strains correspond to the published genome: GenBank CP002487.1 [35]. *Escherichia coli* strains Top10 (Invitrogen) and S17-1 λ*pir* [113] were used as hosts for the cloning procedures.

Unless otherwise specified, bacteria were grown at 37°C with aeration (orbital shaking 220 rpm) in Lennox Broth (LB: 10 g/L BD Tryptone, 5 g/L BD Yeast Extract, 5 g/L NaCl), LBO 10 g/L BD Tryptone, 5 g/L BD Yeast Extract) or in M9 minimal medium [114], prepared with M9 Salts, 2 mM MgSO_4_, 0.1 mM CaCl_2_ and 40 mM sodium succinate dibasic hexahydrate (succinate) or 40 mM glycerol + 10 mM Succinate, as sole C-sources (henceforth, media M9+Succ and M9+Gly+Succ). Agar plates were prepared with the same media, solidified with 1.5% BD Bacto agar. When required, 10-100 µM of FeCl_3_ were added to the M9 media.

To seed the M9-derived media of all the experiments, stationary phase pre-cultures were prepared by inoculating isolated colonies into 5 mL LB (in 30 mL Universal glass tubes) and the cultures were incubated for 6-20 hours at 37°C with aeration. Bacteria were harvested by centrifugation, washed once, and Optical Density at 600 nm (OD_600_) was adjusted to 1 with the minimal medium used for the cultures, or with 1 X Phosphate-Buffered Saline (PBS) to generate a standardised inoculum. Subsequently, bacteria were grown aerobically in conical flasks (topped with aluminium foil) or in Greiner 50 mL plastic tubes (with lids slightly open to allow gaseous exchange). Washed bacteria from the pre-cultures were inoculated as a 1:100 dilution to give a starting OD_600_ of 0.01 (∼10^7^ CFU/mL), in a final medium volume corresponding to 10% of the flask/tube capacity, to ensure optimal oxygenation by shaking.

For growth curves in 96-well microplates (Greiner #655180), bacteria grown beforehand for ∼6 hours in LB, were washed with the minimal medium used for the cultures, or with PBS, and inoculated to give a starting OD_600_ of 0.01 in 200 µL of medium *per* well. The microplates were incubated at 37°C with orbital shaking (500 rpm), in a FLUOstar Omega plate reader (BMG Labtech), and the OD_600_ was monitored every 15-30 min, using the appropriate growth medium as blank.

When required, antibiotics were added as follows: 50 µg/mL kanamycin monosulfate (Km), 100 µg/mL Ampicillin sodium (Ap), 25 µg/mL tetracycline hydrochloride (Tc), 20 µg/mL gentamicin sulfate (Gm) and 25 µg/mL chloramphenicol (Cm).

For strains carrying the *tetR-P_tetA_* module, the *P_tetA_* promoter was induced by adding 500 ng/mL of anhydrotetracycline hydrochloride (AHT, from a 1 mg/mL stock solubilised in methanol). The same volume of methanol was added to the mock-induced cultures. To stimulate expression of genes controlled by the *P_BAD_* promoter (*e.g.* in plasmid pWRG99), 0.2 % L-(+)-arabinose was added to the culture. For the strains carrying the plasmid pSW-2 [115], the *P_m_* promoter was induced by adding 1 mM *m*-toluate (500 mM stock titrated with NaOH to pH 8.0).

### Bacterial transformation and Tn*5* mutagenesis

Chemically-competent *E. coli* were prepared and transformed as previously published [116]. Electrocompetent cells were prepared with *Salmonella* cultures grown in salt free LBO medium and were electroporated, as described previously [117]. After recovery in LB at 37°C (30°C for temperature-sensitive plasmids) transformation reactions were spread on selective LB agar plates and transformants were obtained after incubation at 30°C or 37°C.

For the Tn*5* transposon mutagenesis, ultra-competent *Salmonella* were prepared from LBO cultures grown at 45°C, as previously reported [117, 118]. The λ*pir-*dependent plasmid pRL27 [119], that encodes the Tn*5* transposase gene (*tnp*) and the mini Tn*5*-*ori*R6K-Km^R^ transposon (Tn*5*), was used to generate the *Salmonella* Tn*5* libraries, as follows: 50 µL of ultra-competent *Salmonella* were electroporated with 500 ng of the non-replicating Tn*5* delivery plasmid pRL27. After 1 hour recovery in LB the transformation reactions containing the Tn*5*-carrying *Salmonella* were washed in PBS or minimal media and 1% of transformations was spread on LB Km plates to estimate the size of the resulting Tn*5* library. The remainder of the Tn*5* libraries were stored for further experiments.

### Cloning procedures

Enzymes, buffer and kits used are listed in Supplementary Resource Table S1. DNA manipulations were carried out according to standard protocols [114]. DNA fragment were purified from enzymatic reactions or from agarose gel using the Bioline ISOLATE II PCR and Gel Kit. Plasmids were extracted with the Bioline ISOLATE II Plasmid Mini Kit. Genomic DNA (gDNA) was isolated from 0.5-1 mL of stationary phase cultures with the Zymo Quick DNA Universal Kit.

For PCR, DNA was amplified with Phusion High Fidelity DNA polymerase, template DNA and 0.5 µM primers, in the presence of 3 % Dimethyl Sulfoxide and 1 M betaine. For plasmid/strain verifications by Sanger sequencing, PCR reactions were carried out from bacterial colonies with MyTaq Red Mix 2 X and PCR fragment were Sanger sequenced with the appropriate primers (Lightrun service, Eurofins Genomics).

DNA digestion/ligation procedures or the restriction-free “PCR cloning” technique [120] were used to insert DNA fragments into the plasmids pEMG [115], pJV300 (pP_L_)[67] and pXG10-SF [121]. *E. coli* Top10 was used as host for the construction of pXG10-SF and pP_L_ derived plasmids, while S17-1 λ*pir* was used for pEMG derivatives.

The construction of each plasmid is detailed in Supplementary Resource Table S1. For complementation experiments, the genes of interest (including their native ribosome binding site) were PCR-amplified from 4/74 gDNA and were cloned between the NsiI and XbaI sites of the low copy plasmid pXG10-SF, resulting in plasmids p*hfq* (pNAW45), p*rpoS* (pNAW95), p*iraP* (pNAW98), p*fliST* (pNAW94), p*rbsR* (pNAW93), p*iscR* (pNAW96) and p*cspC* (pNAW92). For the construction of the p*iscR^3CA^* plasmid (pNAW97, carrying three Cys->Ala substitution at positions 92, 98 and 104 in IscR), two fragments carrying the appropriate mutations were first amplified with primer pairs NW_461/NW_467 and NW_462/NW_466 and the resulting amplicons were fused by overlap extension PCR [122] before insertion into pXG10-SF. In all these plasmids, the genes of interest were under the control of the strong constitutive promoter *P_L tetO-1_* [123]. For the construction of p*pnp* (pNAW256), the *pnp* gene and its promoter region (including *sraG*) were amplified and inserted between the XhoI and XbaI sites of pXG10-SF. In the resulting plasmid, *pnp* expression was controlled by its native promoter. For the construction of p*oxyS* (pP_L_-*oxyS*, pNAW255), *oxyS* was PCR amplified and inserted by PCR cloning downstream of constitutive *P_L lacO-1_* promoter of the pP_L_ vector, as described earlier [124]. The plasmid-borne translational fusion *yobF*::*_sf_gfp* (p*yobF*::*_sf_gfp*, pNAW258) was constructed by cloning the 5’-UTR of *yobF* and its 31 first codons in frame with the *_sf_gfp* gene of pXG10-SF, as previously described [121].

Plasmid site-directed mutagenesis [125] was used to introduce the CC->GG mutation in the pP_L_-*oxyS*: complementary primers NW_1022 and NW_1023, carrying the mutations were annealed and elongated by PCR for 10 cycles. Fifty nanograms of plasmid p*oxyS* (pP_L_-*oxyS*, pNAW255) were added to the reaction and the PCR reaction was resumed for 25 cycles. After DpnI treatment and transformation in *E. coli* Top10, the mutated plasmids pP_L_-*oxyS^GG^* (pNAW259) was obtained.

For the construction of the empty vector pNAW125 (pXG10-SF lacking the *lacZ^186^::_sf_gfp* fragment), a 3.5 kb fragment was PCR-amplified from pXG10-SF with the primer pair NW_348/NW_565, the resulting fragment was digested with NsiI and self-ligated.

All the pXG10-SF, pEMG and pP_L_ derived plasmids were verified by Sanger sequencing: primers pXG10_R2 and pZE-CAT were used for plasmid pXG10-SF, primers M13_-40_long and M13_Rev_long for plasmid pEMG and primer pZE-A for plasmid pP_L_.

### Genome editing techniques

The λ *red* recombination methodology was used to insert or delete genes in the *Salmonella* chromosome, using the heat-inducible λ *red* plasmid pSIM5-*tet* [127]. *Salmonella* carrying the pSIM5-*tet* were grown in LBO at 30°C and electrocompetent bacteria were prepared after a 15 minute heat shock at 42°C, as previously described [117, 128]. PCR fragments carrying a resistance gene were PCR-amplified from the template plasmids pKD4, pKD3, pNAW52, pNAW55 or pNAW62 [117, 126]. Electrocompetent bacteria (50 µL) were transformed with 500-3000 ng of the PCR fragments, and recombinants were selected on LB agar plates that contained the appropriate antibiotics.

The deletions/insertions linked to a selective marker and the Tn*5* insertions were transduced to *S.* Typhimurium strains using the P22 HT *105/1 int-201* (P22 HT) transducing phage, as previously reported [129, 130].

When required, antibiotic resistance cassettes flanked by FLP recognition target sites (*frt*), were removed from the *Salmonella* chromosome with the FLP recombinase-expressing plasmid pCP20 [131]. Subsequently, the temperature sensitive plasmid pCP20 plasmid was eliminated by a single passage at 42°C.

The construction of each strain is detailed in Supplementary Resource Table S1. The marked mutations Δ *S*::*aph*, Δ *hfq*::*aph,* Δ *rybB*::*aph,* Δ:: *cat* (*pnp-539*) and Δ*arcA*::*aph* were transduced from published 4/74 and SL1344 derivative strains. For the transduction of the Δ *q*::*aph* mutation, the donor strain JH3584 was first complemented with the plasmid p*hfq* (pNAW45), because Hfq is required for P22 transduction [67].

The 4/74 *tetR-P_tetA_*-*dctA* strain was constructed according to the principle described by Schulte and colleagues [132]. The promoter and the 5’-UTR of *dctA* (coordinates 3812883-3812972) were replaced by a *frt-aph-frt-tetR-P_tetA_* module from plasmid pNAW55. In the resulting strain, *dctA* has the *tetA* ribosome binding site and is controlled by the AHT-inducible *P_tetA_* promoter.

To measure *rpoS* expression, a chromosomal transcriptional *rpoS-gfp*^+^ fusion was constructed as follows: the *gfp^+^-frt-aph-frt* module of pNAW52 (including the *gfp^+^* SD) was inserted after the main *rpoS* transcription start site (TSS), previously mapped at the coordinate 3089613 on the 4/74 chromosome [133].

To measure *dctA* expression, a chromosomal transcriptional/translational *dctA*::*_sf_gfp* fusion was constructed: the flexible amino acid linker GSAGSAAGSGEF and the sequence encoding for superfolder GFP (*_sf_gfp*) were fused to the *dctA* C-term, using a *_sf_gfp-frt-aph-frt* module amplified from pNAW62 [117].

For complementation in *oxyS* and *rpoS* null mutants, a single copy of these genes was inserted in a non-transcribed chromosomal locus in two steps: first, an antibiotic cassette was inserted upstream of the *oxyS* or *rpoS* promoters, coordinates 4364521 and 3089777, respectively. Then, the *aph-oxyS* and the *cat*-*rpoS* modules were PCR-amplified from the resulting strains and inserted into the non-transcribed pseudogene *STM474_1565* (between coordinates 1585970-1586170 on the 4/74 chromosome) [117]. The *STM474_1565* gene is also known as *STM1553* or *SL1344_1483*.

For scarless transfer of *rrsA^mut^* and *rrsH^mut^* mutations located in the 16S rRNA genes, the two step λ *red* recombination-based methodology described by Blank and colleagues was used [134]. The *rrsA* and *rrsH* genes (including their promoters) were replaced by a I-*Sce*I-*aph* module, amplified from plasmid pKD4-I-*Sce*I [129]. The mutations were transduced in 4/74 WT, yielding the strains Δ*rrsA*::(I-*Sce*I-*aph*) and Δ*rrsA*::(I-*Sce*I-*aph*). The two strains were transformed with the *red* plasmid pWRG99, expressing the I-SceI nuclease in the presence of AHT [134]. Electro-competent cells were prepared with the two *Salmonella*+pWRG99 strains in the presence of arabinose, as described above. Competent bacteria were electroporated with 2 µg of the *rrsA^mut^* or the *rrsH^mut^* fragments, obtained by PCR from spontaneous mutants SNW245 (*rrsA^mut^*) and SNW246 (*rrsH^mut^*). Replacement of the I-*Sce*I-*aph* module by the corresponding *rrs* mutated PCR fragment was selected on LB agar supplemented with Ap and AHT at 30°C. The *rrsA^mut^* or *rrsH^mut^* insertions were confirmed by PCR and Sanger sequencing and the temperature-sensitive pWRG99 plasmid was eliminated by a passage at 42°C.

For other scarless genome editing procedures, the pEMG-based allele exchange technique [115] was used as previously described [129, 135]. For the transfer of the *dctA^mut1^*, *dctA^mut2^* and *oxyR^mut^* mutations, fragments encompassing the mutations and the ∼500 bp flanking regions were PCR-amplified from the corresponding spontaneous Succ^+^ mutants. For the construction of the *oxyS^GG^* mutant, two fragments flanking the mutations were PCR amplified with primer pairs NW_1021/NW_1022 and NW_1023/NW_1024. The two amplicons carrying the CC->GG mutation on one of their extremities were fused by overlap extension PCR. All the PCR fragments carrying the mutations were inserted by digestion/ligation between the EcoRI and BamHI sites of the suicide plasmid pEMG and *E. coli* S17-1 λ*pir* was transformed with the resulting ligation reactions. The resulting suicide plasmids were mobilised into the recipient *Salmonella* by conjugation and recombinants were selected on M9-glucose agar plates containing Km. Merodiploid resolution was carried out with the pSW-2 plasmid, as previously described [129]. The relevant mutations were confirmed in Km^S^ candidates by PCR and Sanger sequencing. Finally, the unstable pSW-2 plasmid was eliminated by 2-3 passages on LB.

### Experimental evolution to select Succ^+^ mutants

A 4/74 strain with an additional *rpoS* copy inserted in the *STM474_1565* pseudogene (strain *rpoS^2X^,* SNW226, Cm^R^) was transformed with pRL27 and ten libraries of approximately 10,000 Tn*5* mutants were grown aerobically in 25 mL of M9+Succ containing Cm and Km in ten 250 mL conical flasks. After 48 hours incubation at 37°C, the cultures were spread on LB + Km plates and isolated colonies were passaged twice on LB agar. Growth on M9+Succ plates was assessed for >10 isolates *per* library. The Tn*5* insertions from fast-growing colonies (Succ^+^ phenotype) were P22-transduced into 4/74 WT, and the Succ^+^ status of the transductants was verified.

For the isolation of Succ^+^ spontaneous 4/74 mutants, bacteria obtained from stationary phase LB cultures were washed with PBS and the OD_600_ was adjusted to 0.1 (∼10^8^ CFU/mL). Approximately 10^7^ CFU (100 µL) were spread on M9+Succ agar and the plates were incubated at 37°C, until Succ^+^ large colonies were visible (3-4 days). Alternatively, spontaneous mutants were obtained from liquid M9+Succ cultures (25 mL), inoculated with ∼100 *Salmonella.* The cultures were grown aerobically at 37°C, until substantial growth was observed (typically after 3 days incubation) and the cultures were spread on LB agar plates. All the presumed Succ^+^ spontaneous mutants were passaged twice on LB plates before confirming the Succ^+^ phenotype on M9+Succ agar plates. The *rpoS* positive status (*rpoS*^+^) of each mutant was tested by phenotypic assays (see below) and confirmed by PCR and Sanger sequencing, using primers NW_403, NW_252 and NW_252.

The genomes of a collection of *rpoS*^+^ Succ^+^ mutants were sequenced by the Illumina whole genome sequencing service of MicrobesNG (Birmingham, UK). Mutations were identified using the VarCap workflow [136] available on Galaxy (http://galaxy.csb.univie.ac.at:8080), using the published 4/74 genome as reference. The identified mutations were confirmed by PCR and Sanger sequencing, and were transferred into 4/74 WT using the two scarless genome editing techniques described above. After transfer, the Succ^+^ phenotype of all genome-edited mutants was confirmed.

### Phenotypic characterisation of the Succ^+^ mutants

Phenotypes linked to the *rpoS* status were tested for each of the Succ^+^ Tn*5* or Succ^+^ spontaneous mutants. The RpoS-dependent catalase activity was assessed with hydrogen peroxide directly on colonies or with stationary phase LB cultures, as described earlier [137, 138]. The RpoS-dependent RDAR (red, dry and rough) morphotype [87] was tested by adding 2 µL of a stationary phase LB cultures on LBO agar plates containing 40 μ mL of Congo Red. The RDAR morphotype was observed after at least 3 days of incubation at room temperature.

### Mapping of Tn*5* insertion sites

The Tn*5* insertion sites of the Succ^+^ mutants were mapped by an arbitrary PCR approach [139]. For each Tn*5* mutant, the arbitrary PCRs were carried out directly from colonies with the primer pair NW_319/NW_320 (0.5 µM each) and MyTaq Red Mix 2 X in a final volume of 20 µL. The arbitrary PCR conditions were: 95°C 120 sec; 6 X [95°C 15 sec 30°C; 30 sec; 72°C 90 sec]; 30 X [95°C 15 sec; 50°C 30 sec; 72°C 90 sec]; 72°C 300 sec; 4°C. The amplicons were purified on column and eluted in 20 µL of water. For the nested PCRs, 2 µL of the arbitrary PCR products were used as template and mixed with primer pair NW_318/NW_321 (0.5 µM each) and MyTaq Red Mix 2 X in a final volume of 40 µL. The second PCR conditions were: 95°C 120 sec; 30 X [95°C 15 sec; 50°C 30 sec; 72°C 90 sec]; 72°C 300 sec. The PCR products were separated by electrophoresis on a 1 % agarose gel containing Midori Green for DNA UV-visualization. The most prominent DNA bands were excised, the DNA was purified and Sanger-sequenced with primer NW_318. Insertions were mapped by BLAST, using the 4/74 genome as reference.

### Quantification of GFP fluorescence intensity

Strains carrying the chromosomal fusion *rpoS-gfp*^+^ or the plasmid-borne *yobF*::*_sf_gfp* translational fusion were grown in the indicated conditions in biological triplicates and bacteria were harvested by centrifugation and re-suspended in the same volume of PBS. The GFP signal was measured with a FLUOStar Omega plate reader (BMG Labtech) with 200 µL of bacterial suspension *per* well in black microplates (Greiner #655090). For each strain, the PBS background fluorescence was subtracted from the GFP signal (in arbitrary unit [a.u.]). The fluorescence values were divided by the OD_600_ of the cell suspensions. The fluorescence background of a WT unlabeled strain (carrying the empty plasmid pNAW125, when required) was measured similarly, and was subtracted from the fluorescence signal of the GFP-labelled strain. For each strain the GFP fluorescence intensity is represented as absolute values (GFP fluorescence intensity/OD_600_ [a.u.]) or as a relative GFP fluorescence intensity (in %).

To measure the activity of the *dctA*::*_sf_gfp* chromosomal fusion, bacteria were grown in M9+Gly+Succ to OD_600_ 0.5-1. Cells were harvested and re-suspended in PBS, prior to fixation with 4% paraformaldehyde and washes with PBS [140]. To measure the GFP fluorescence intensity in all the strains carrying *dctA*::*_sf_gfp*, the IntelliCyt iQue® Screener PLUS (Sartorius) was used. To quantify the *dctA*::*_sf_gfp* activity more precisely in the WT, *dctA^mut1^*, *dctA^mut2^,* Δ*iscR* and Δ*rpoS* genetic backgrounds, bacteria were grown in biological triplicates and were fixed with paraformaldehyde. The FITC-H GFP fluorescence intensity (median of the population) was measured using a FACSCanto^TM^ II flow cytometer (BD Biosciences). The fluorescence background of a WT unlabelled strain was subtracted from the fluorescence intensity of each *dctA*::*_sf_gfp* carrying strain. The data are represented as the GFP fluorescence intensity of each mutant, relative to the intensity of the WT isogenic strain (%). All flow cytometry data were analysed using the FlowJo^TM^ software (BD Biosciences).

For fluorescence microscopy, the *dctA*::*_sf_gfp*-carrying strains were grown in M9+Succ or M9+Gly+Succ and bacteria were immobilised in PBS solidified with 0.75% low melting point agarose. Pictures were taken with the EVOS FL cell imaging system (Thermo Fisher), as previously described [124].

### RpoS detection by Western blotting

The strains of interest were grown in the indicated condition, and bacteria (∼10^9^ CFU, estimated by OD_600_) were pelleted by centrifugation and stored at −80°C. Bacteria were re-suspended in 100 µL of PBS and 10 µL of the cell suspensions were mixed with 990 µL of PBS to measure the OD_600_ 1/100 of each suspension. Bacteria were lysed by adding 100 µL Laemmli Buffer 2 X [120 mM Tris-HCl pH 6.8, 4% (wt/vol) SDS, 20% (vol/vol) glycerol, Bromophenol blue 0.02% (wt/vol)] and 10 μ β-mercaptoethanol (5% vol/vol final). The lysates were boiled for 15 min, chilled on ice for 1 min and spun down for 5 min at 4 °C (14,000 rpm). Bacterial extracts were separated by SDS polyacrylamide gel electrophoresis and the proteins RpoS and DNaK were detected by Western Blotting, as described earlier [128]. The volume of protein extract loaded (∼10 µL) on the SDS 10% polyacrylamide gel was normalised by the OD_600_ of the bacterial suspensions prior to lysis.

Catalogue numbers of all antibodies are listed in Supplementary Resource Table S1. The primary antibodies, Anti-*E. coli* RNA Sigma S Antibody (diluted 1:5,000) and anti-DnaK mAb 8E2/2 (diluted 1:10,000), were used for the detection of RpoS and DnaK (loading control), respectively. For detection, the secondary antibody Goat anti-mouse IgG (H + L)-HRP (diluted 1:2,500) and the Pierce ECL Western blotting substrate were used. The chemiluminescent reaction was detected with the ImageQuant LAS 4000 imaging system (GE Healthcare Life Sciences).

### OxyS detection by Northern blotting

The WT, Δ*oxyR* and *oxyR* strains were grown in 25 mL of LB to OD_600_ = 1 and the cultures were split in two 10 mL subcultures. For each strain, hydrogen peroxide (H_2_O_2_, 2 mM) was added to one of the subcultures. After 30 min at 37 °C, cellular RNA transcription and degradation processes were stopped by adding 4 mL of ice-cold STOP solution (95% ethanol + 5% acid phenol) to the 10 mL cultures. After a 30 min incubation on ice, bacteria were pelleted by centrifugation and total RNA was extracted with Trizol, as described previously [124].

Probe synthesis and OxyS sRNA detection by Digoxigenin (DIG)-based Northern blotting were carried out with the DIG Northern Starter Kit, according to the DIG Application Manual for Filter Hybridization (Roche) and a previous study [124]. Briefly, heat-denatured RNA (2.5 µg) was separated on an 8.3 M urea, 7% polyacrylamide gel in TBE 1X. RNA was transferred to a positively charged nylon membrane with the Bio-Rad Semi Dry transfer system (#170-3940). RNA was UV-crosslinked to the membrane before hybridization with the DIG-anti-OxyS probe in DIG Easy Hyb buffer at 68°C for 20 hours. The membrane was washed and the OxyS transcripts were detected using the Anti-Digoxigenin antibody and the CDP-*Star* substrate. Finally, the chemiluminescent reaction was visualised using the ImageQuant LAS 4000 imager. After OxyS detection, the membrane was stripped and re-probed with the DIG-anti-5S probe to detect the 5S ribosomal RNA, used as a loading control. The ssRNA DIG-labelled probes DIG-anti-OxyS and DIG-anti-5S were synthesised with the T7 polymerase and DNA templates obtained by PCR with template 4/74 gDNA and primer pairs NW_485/NW_485 and DH58/DH59, respectively.

### Quantification and statistical analysis

Numerical data were plotted and analysed using Microsoft Excel (version 16.46). Data are presented as the mean of three to six biological replicates ± standard deviation, as indicated in the figures. The unpaired *t-*test was used to compare the groups and statistical significance is indicated on the figures. *P* values (two-tail) are reported using the following criteria: 0.0001 to 0.001 = ***, 0.001 to 0.01 = **, 0.01 to 0.05 = *, ≥ 0.05 = NS

## Supporting information

Supplementary Resource Table S1

## Acknowledgements

We are very grateful to Aoife Colgan for the construction of several mutants, to Stefano Marzi and Laurent Aussel for helpful advice, and to Paul Loughnane for his expert technical assistance. We also thank Celeste Peterson and Susan Gottesman for sharing preliminary data about the *rbs*-driven regulation of RpoS. Finally, we thank all the present and former members of the Hinton Lab for helpful and productive discussions (in the AJ and elsewhere).

## Supplementary Figures

**Figure S1.**
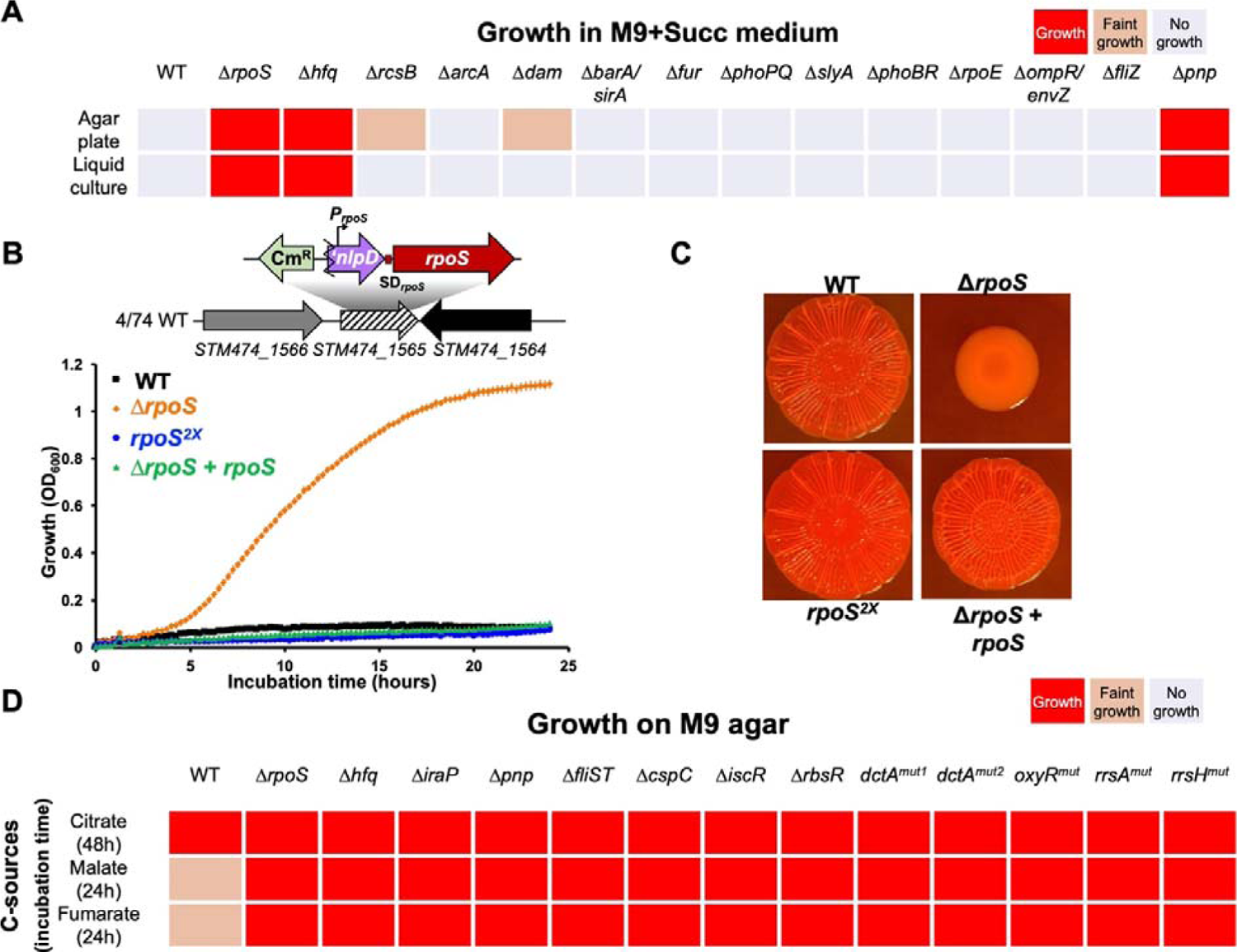
Growth assessment of *Salmonella* mutants with diverse C-sources involving a chromosomal construct to complement the *rpoS* mutation. (**A**) the growth of a collection of *S*. Typhimurium 4/74 mutants lacking regulatory proteins was assessed on solidified M9+Succ agar and in liquid M9+Succ medium (in microplates), revealing the Δ*pnp* (JH3649). (**B**&**C**) Chromosomal complementation of the Δ*rpoS* mutation: a copy of *rpoS* (including its native promoter, bent arrow), linked to the *cat* Cm resistance gene was inserted in the non-transcribed pseudogene *STM474_1565* of 4/74 WT (strain 4/74 *rpoS^2X^*, SNW226) and of Δ*rpoS* ( *rpoS + rpoS*, JH4160). The *STM474_1565* gene is also known as *STM1553* or *SL1344_1483*. RpoS-dependent phenotypes were assessed for each strain: growth was tested in M9+Succ (**B**) and RDAR phenotype was tested on Congo Red agar plates, confirming the Succ^-^ RDAR^+^ of the complemented strain Δ*rpoS + rpoS*. (**D**) The growth of the novel Succ^+^ mutants identified (presented in Fig 3) was tested on solidified M9 minimal medium supplemented with 40 mM citrate, malate or fumarate. The growth was assessed with biological triplicates after the indicated incubation time (37°C) and the growth of each mutant was compared to 4/74 WT (Succ^-^) and Δ*rpoS* (Succ^+^).

**Figure S2.**
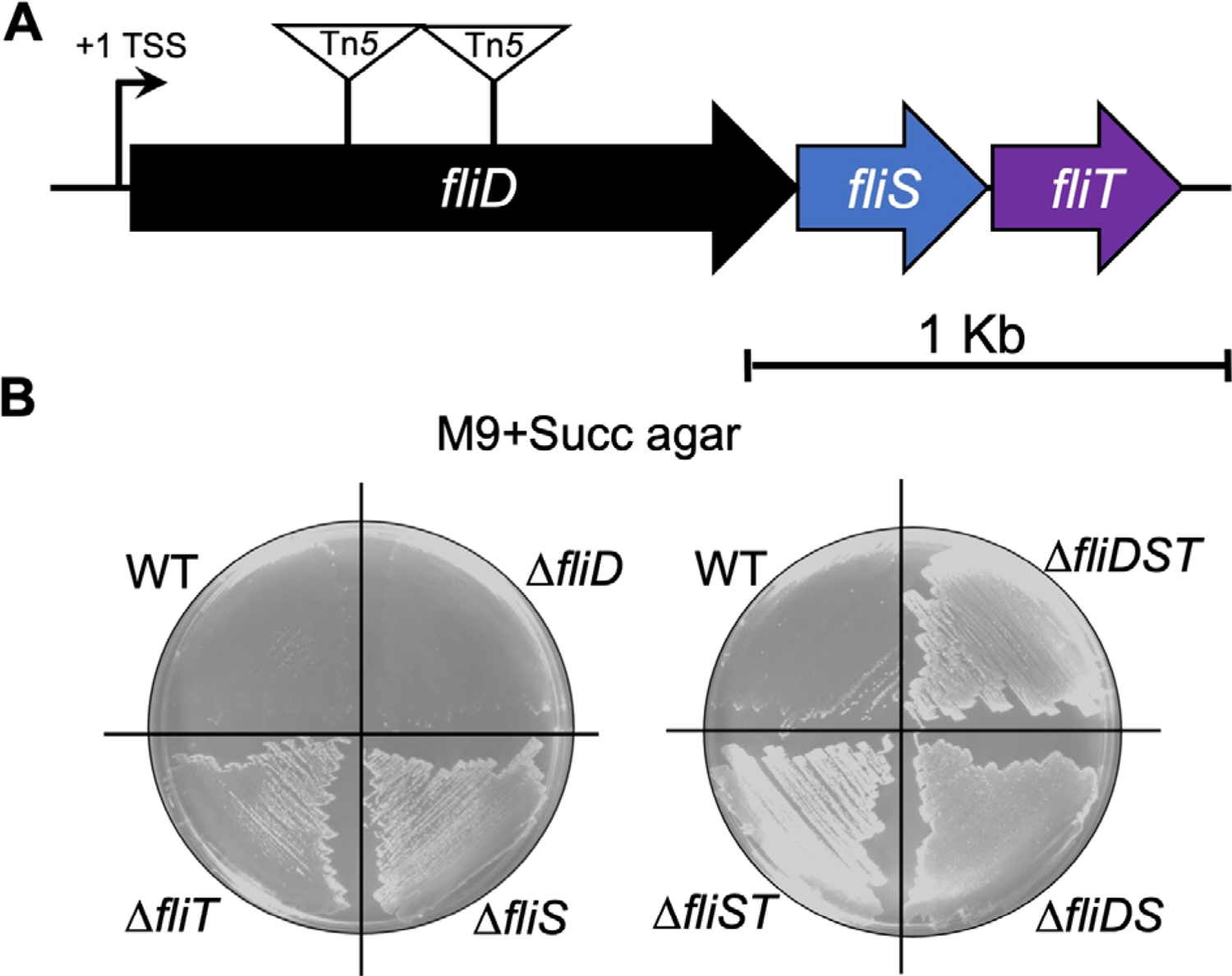
Genetic dissection of the *fliDST* operon reveals that *fliS* and *fliT* can inhibit succinate utilisation. (**A**) Schematic representation of the *fliDST* operon. The transcription start site (+1 TSS) and the two Tn*5* transposon insertions causing Succ^+^ phenotype are depicted (Table 1). (**B**) The inactivation of the flagellar chaperones FliS and FliT stimulates *Salmonella* growth with succinate. The growth of 4/74 WT and of mutants Δ*fliD* (SNW278), Δ*fliS* (SNW280), Δ*fliT* (SNW282), Δ*fliDST* (SNW284), Δ*fliDS* (SNW286), Δ*fliST* (SNW288) was assessed on M9+Succ agar plates after 48 hours of incubation at 37°C.

**Figure S3.**
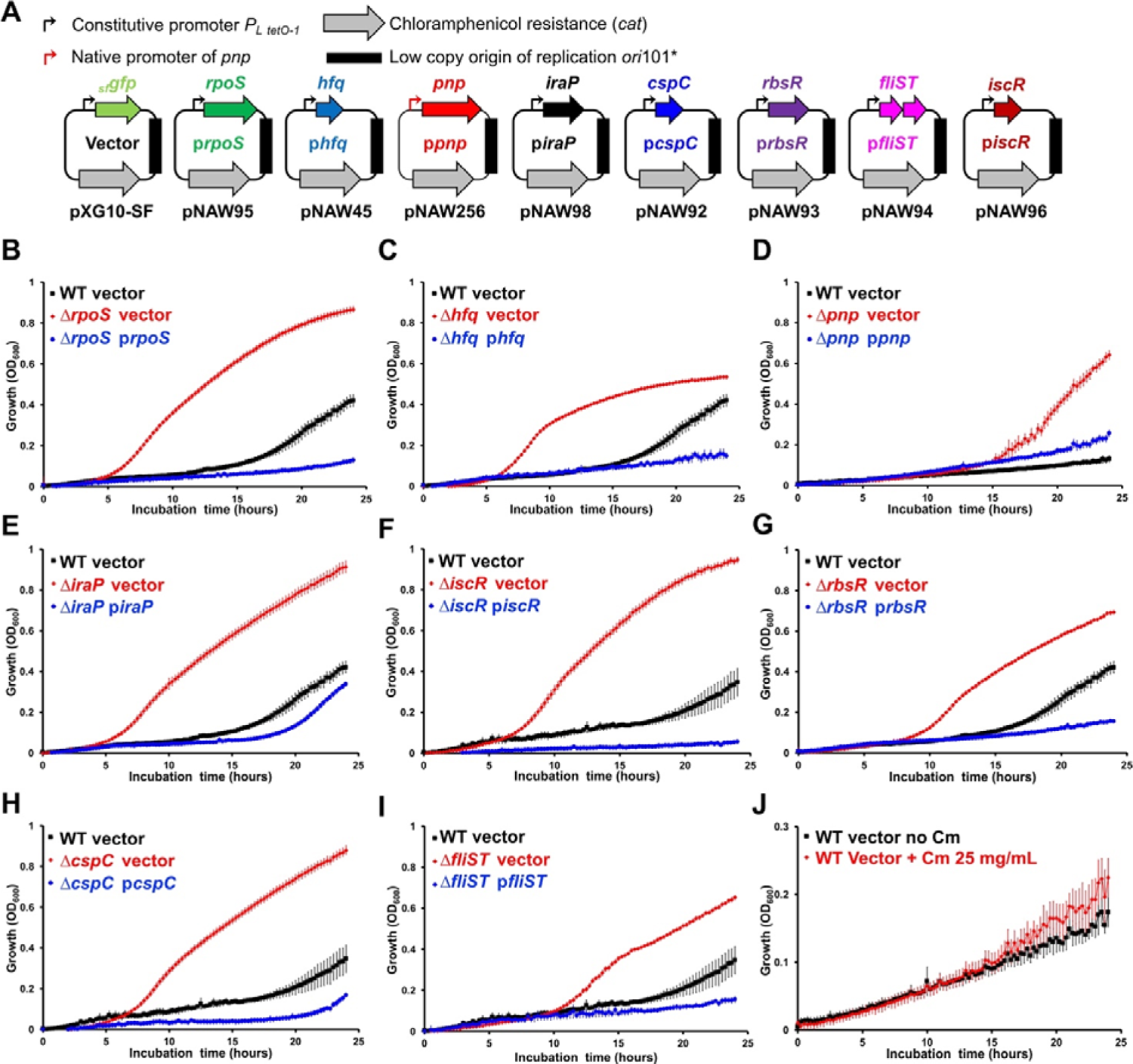
Genetic complementation of eight Succ^+^ regulatory mutations. (**A**) The plasmids used for the complementation experiments are depicted and were constructed (Methods) using the backbone of pXG10-SF [121], a low-copy plasmid encoding for the *cat* resistance gene and carrying the *ori*101* replicon. Each plasmid carries the gene(s) of interest under the control of the strong constitutive promoter *P_L tetO-1_* [123], except for *pnp*, that is controlled by its native promoter. For each growth curve (**B-I**), the plasmid pXG10-SF (“vector”) expressing the *lacZ^186^*::*_sf_gfp* fusion was used as a negative control. The strains 4/74 WT, Δ*S* (JH3674), Δ*q* (JH3584), Δ(JH3649), Δ(SNW188), Δ *cR* (SNW184), Δ*R* (SNW294), Δ*C* (SNW292) and Δ*iST* (SNW288) carrying the indicated plasmids were grown in M9+Succ, supplemented with 25 µg/mL Cm. (**J**) The presence of Cm (25 µg/mL) in M9+Succ stimulates mildly the growth of 4/74 WT carrying pXG10-SF. The growth curves were carried out with 6 replicates in 96-well plates.

**Figure S4.**
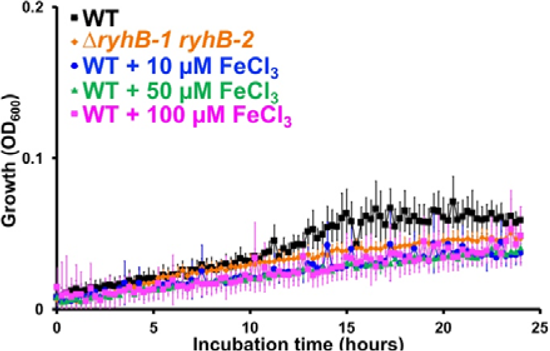
*Salmonella* growth with succinate is not stimulated by iron supplementation or the inactivation of sRNAs RyhB-1 and RyhB-2. Strains 4/74 and Δ*ryhB-1 ryhB-2* (Δ*ryhB-1/2*, JH4390) were grown in M9+Succ medium supplemented or not with iron (FeCl_3_) at the indicated concentration. The growth curves were carried out with 6 replicates in 96-well plates in the indicated medium.

**Figure S5.**
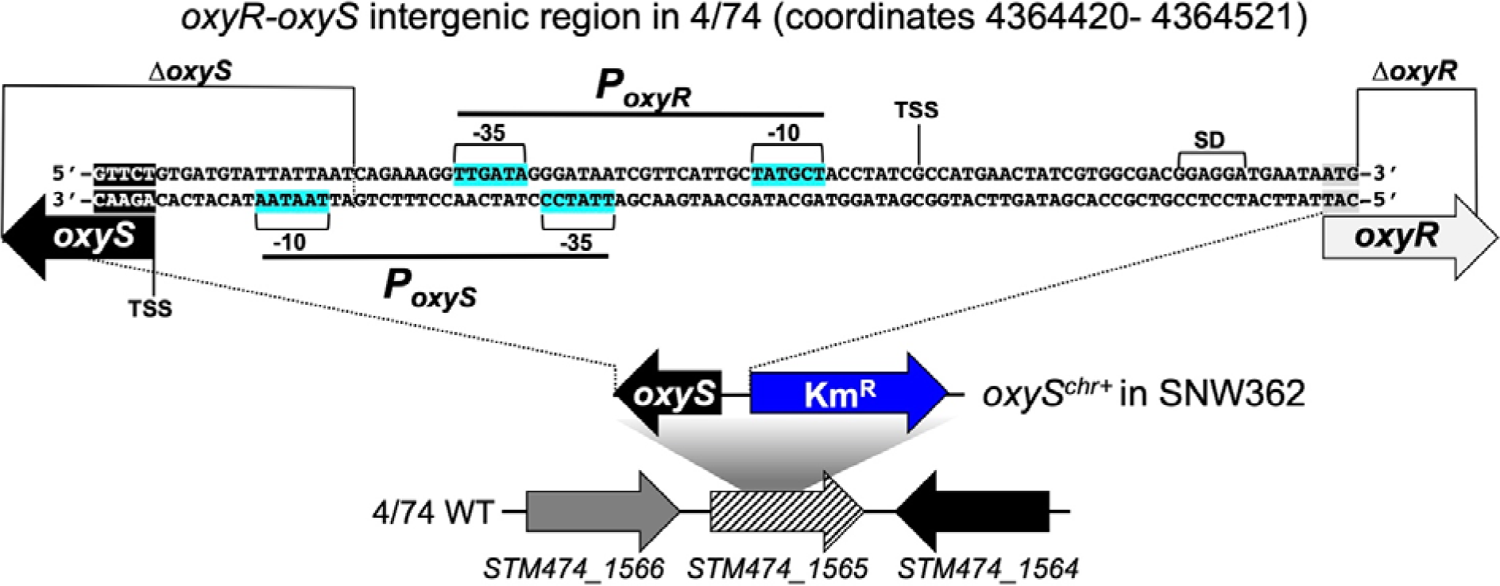
Detailed schematic representation of the *oxyS-oxyR* intergenic region and of the Δ*oxyS*, Δ*oxyR* and Δ*oxyR^chr+^* constructs. The intergenic region sequence is depicted and the −35 and −10 boxes of the *P_oxyS_* and *P_oxyR_* promoters are highlighted in blue, according to the corresponding locus of *E. coli* K-12 [59]. The Δ *oxyS* and Δ*oxyR* mutation are indicated. The transcription start sites (TSS) are indicated, according to the SalcomMac transcriptomic database [108, 133]. For the complementation of the Δ*oxyS* mutation in strain SNW362 (*oxyS^chr+^*), the *oxyS* gene, its native promoter and a Km^R^ cassette were inserted into the non-transcribed pseudogene *STM474_1565* [117].

**Figure S6.**
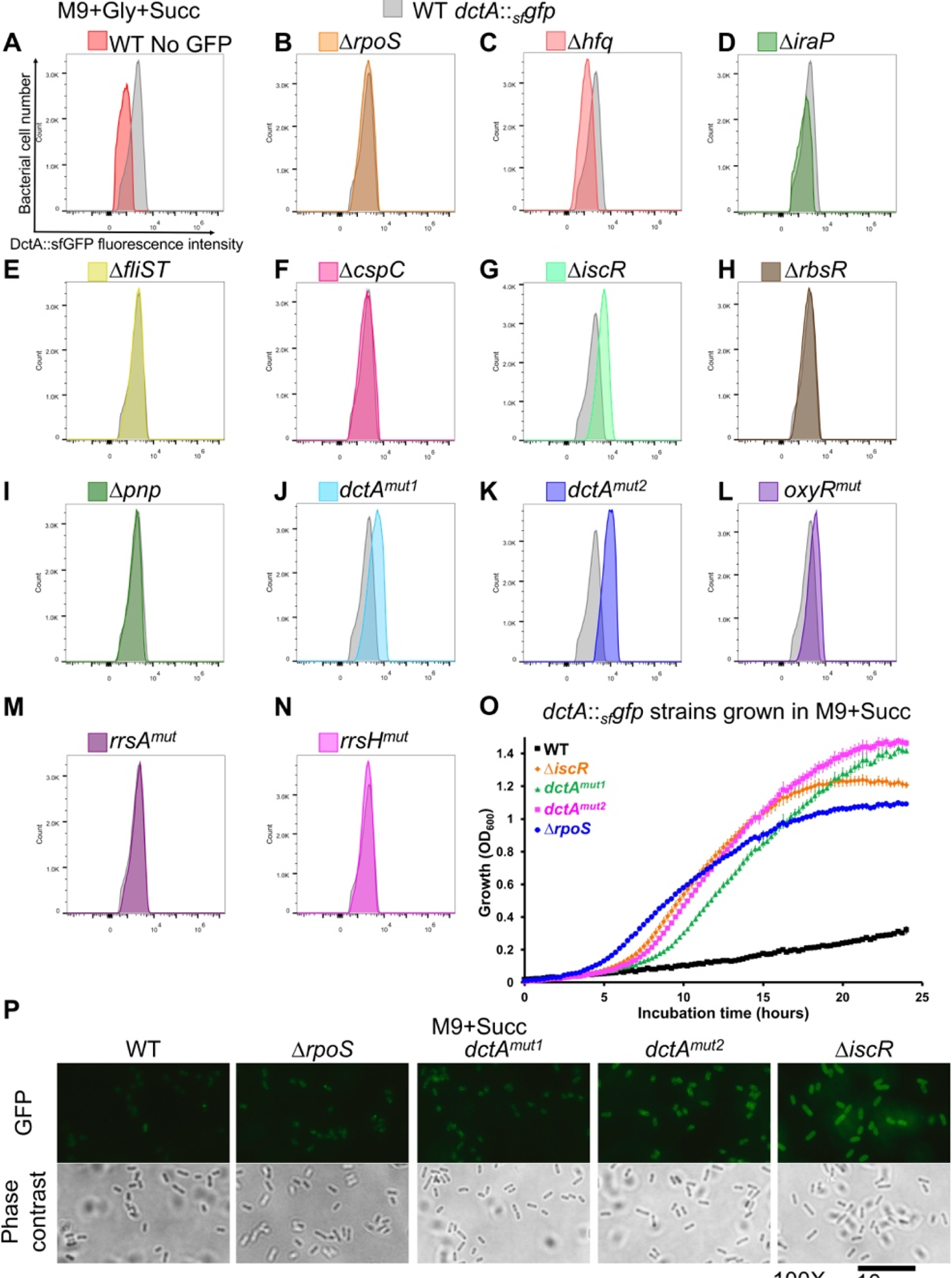
Stimulation of *dctA* expression in the *dctA^mut1^*, *dctA^mut2^* and Δ*iscR* mutants. (**A-N**) Strains carrying the chromosomal transcriptional/translational fusion *dctA*::*_sf_gfp* were grown in M9+Gly+Succ minimal medium (OD_600_∼1) and the GFP fluorescence intensity was measured with the IntelliCyt iQue® Screener PLUS (Sartorius) after bacteria fixation with formaldehyde. The 4/74 WT (untagged strain) was used as a negative control (**A**). Each Succ^+^ mutants carrying *dctA*::*_sf_gfp* was compared with the “WT” strain carrying the same fusion (SNW296, in grey). (**O**) The *dctA*::*_sf_gfp* tagged strain Δ*iscR, dctA^mut1^, dctA^mut2^* and Δ*rpoS* grow fast in M9+Succ in comparison with the *dctA*::*_sf_gfp* tagged WT strain, showing that the fusion of sfGFP to the C-term of DctA does not impede the DctA-driven uptake of succinate. (**P**) The same strains were grown in M9+Succ (OD_600_∼1) and the *dctA*::*_sf_gfp* induction was observed by fluorescence microscopy, as specified in Methods. The *dctA*::*_sf_gfp* tagged Succ^+^ mutants used for these experiments were: Δ*rpoS* (SNW313), Δ*q* (SNW309), Δ*iraP* (SNW423), Δ*iST* (SNW330), Δ *C*(SNW424), Δ*iscR* (SNW329), Δ*rbsR* (SNW425), Δ*pnp* (SNW437), *dctA^mut1^* (SNW310), *dctA^mut2^* (SNW316), *oxyR^mut^* (SNW426), *rrsA^mut^* (SNW374) and *rrsH^mut^* (SNW331).

**Figure S7.**
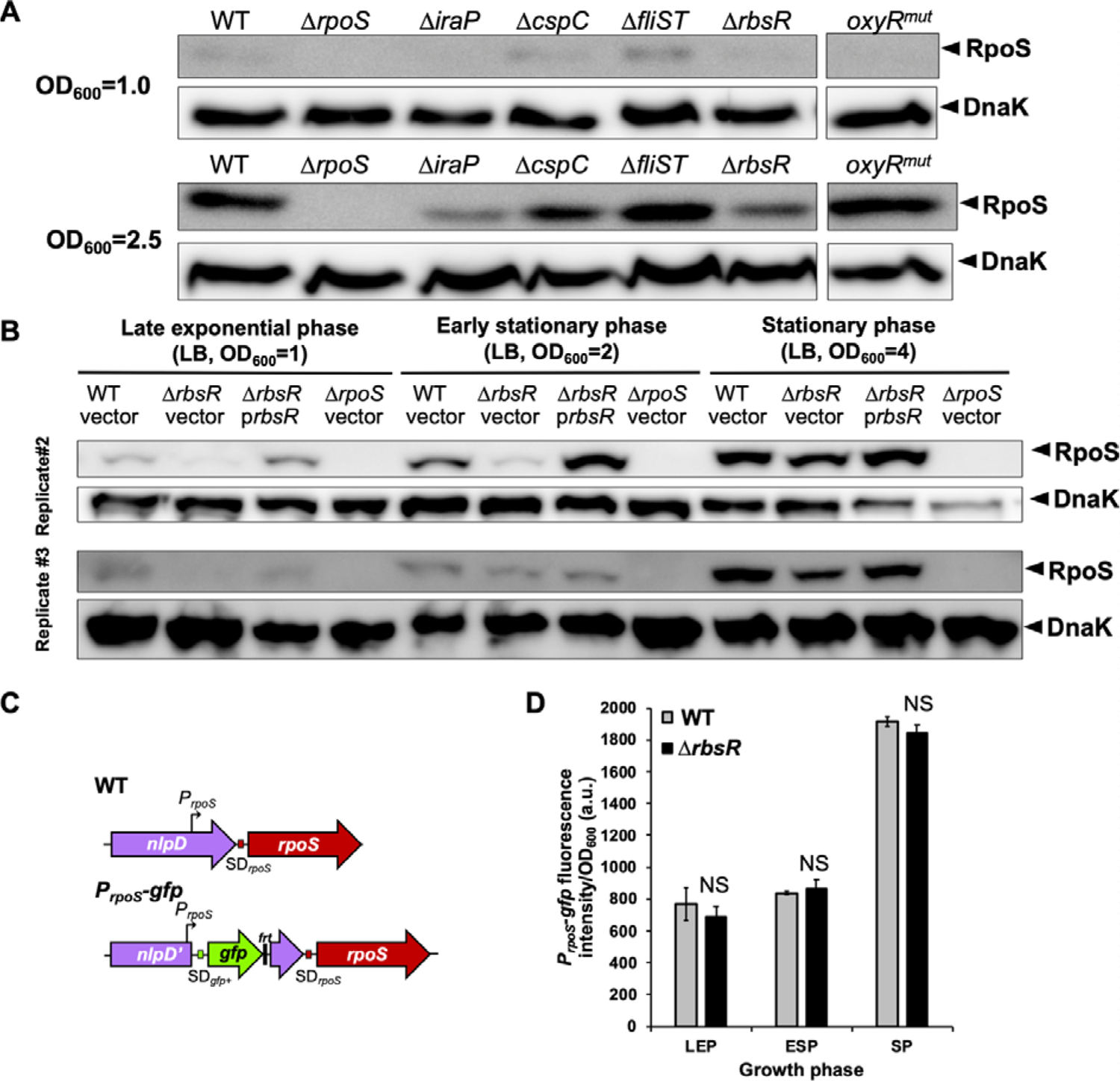
RbsR stimulates *rpoS* expression at the protein level but not at the transcriptional level. (**A**) Western blot detection of RpoS and DnaK (loading control) in 4/74 WT and mutants Δ*rpoS* (JH3674), Δ*iraP* (SNW188), Δ*cspC* (SNW292), Δ*fliST* (SNW288), Δ*rbsR* (SNW294) and *oxyR^mut^* (SNW318) grown in LB to OD_600_ 1 and 2.5. (**B**) Two independent replicates of the Western blot analyses presented in Fig 8A confirmed the down-regulation of *rpoS* in the Δ*rbsR* mutant (see Fig 8A legend). (**C**) Schematic representation of chromosomal *P_rpoS_*-*gfp* transcriptional fusion. The *gfp+* gene and its Shine-Dalgarno (SD) were inserted downstream of the main promoter of *rpoS* (*P_rpoS_*, bent arrow), interrupting the *nlpD* gene. The residual FLP recognition target site sequence is denoted by “*frt*”. The *P_rpoS_*-*gfp* fusion was inserted in 4/74 WT and in Δ*rbsR*, resulting in strain SNW367 and SNW368, respectively. (**D**) The *P_rpoS_*-*gfp* fusion activity was measured in the WT and Δ*rbsR* genetic background in bacteria grown in LB to late exponential phase (LEP, OD_600_∼1), early stationary phase (ESP, OD_600_∼2) and stationary phase (SP, OD_600_∼4). The GFP fluorescence intensity (absolute values) were measured, as specified in Methods. The data are presented as the average of biological triplicates ± standard deviation. The difference of fluorescence intensities between the WT and the Δ*rbsR* strains were not significant (NS) in the three conditions tested, as defined in Methods.

**Figure S8.**
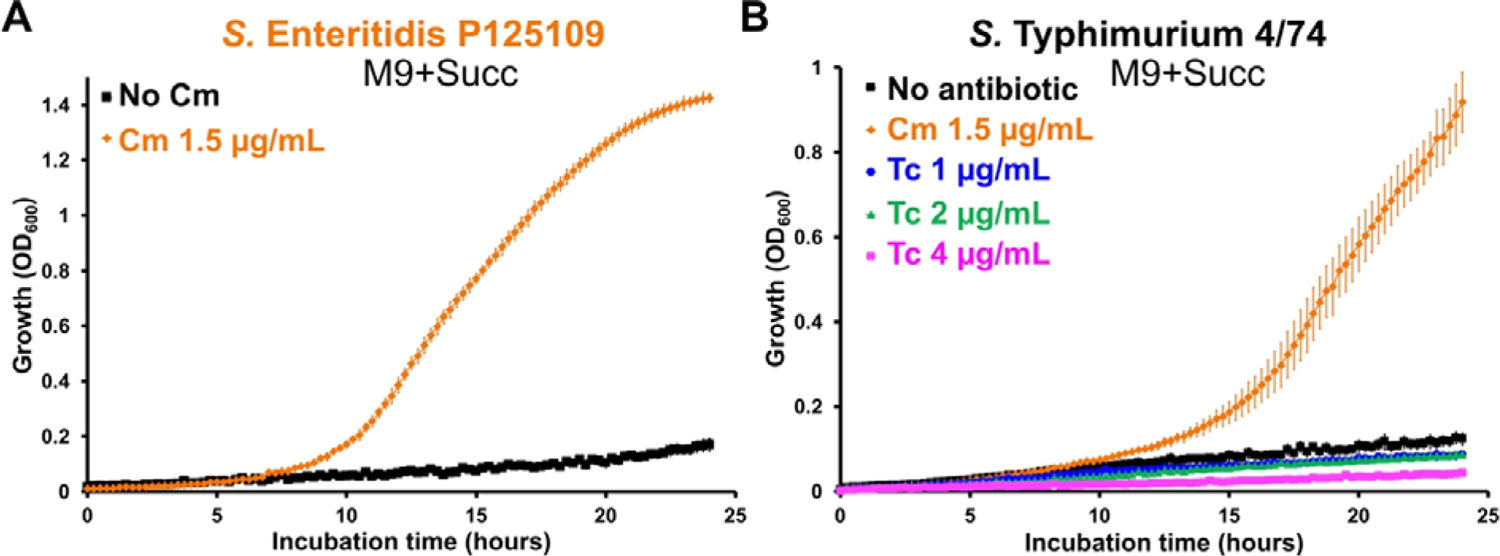
Subinhibitory concentrations of chloramphenicol, but not of tetracycline, stimulate *Salmonella* growth with succinate. (**A**) Low concentration of chloramphenicol (Cm) stimulates the growth of *S.* Enteritidis strain P125109 and of *S.* Typhimurium strain 4/74 with succinate, while tetracycline (Tc) does not affect the growth profile (**B**).

